# *In Toto* imaging of early Enteric Nervous System Development reveals that gut colonization is tied to proliferation downstream of Ret

**DOI:** 10.1101/2022.08.31.506106

**Authors:** Phillip A. Baker, Rodrigo Ibarra-García-Padilla, Akshaya Venkatesh, Eileen W. Singleton, Rosa. A. Uribe

## Abstract

The enteric nervous system (ENS) is a vast intrinsic network of neurons and glia within the gastrointestinal tract and is largely derived from enteric neural crest cells (ENCCs) that emigrate into the gut during vertebrate embryonic development. Study of ENCC migration dynamics and their genetic regulators provides great insights into fundamentals of collective cell migration and nervous system formation, and are a pertinent subject for study due to their relevance to the human congenital disease, Hirschsprung disease (HSCR). For the first time, we performed *in toto* gut imaging and single-cell generation tracing of ENCC migration in WT and a novel *ret* heterozygous background zebrafish (*ret*^*wmr1/+*^) to gain insight into ENCC dynamics *in vivo*. We observed that *ret*^*wmr1/+*^ zebrafish produced fewer ENCCs while localized along the gut, which failed to reach the hindgut, resulting in HSCR-like phenotypes. Specifically, we observed a proliferation dependent migration mechanism, where cell divisions were associated with inter-cell distances and migration speed. Lastly, we detected a premature neuronal differentiation gene expression signature in *ret*^*wmr1/+*^ ENCCs, collectively suggesting that Ret signaling may function to regulate maintenance of a stem-state in ENCCs.

## Introduction

The vertebrate enteric nervous system (ENS) consists of a series of interconnected neurons and glia that form nerve plexuses spanning circumferentially within the muscle walls of the entire gastrointestinal (GI) tract. As the largest component of the peripheral nervous system, the ENS enables the GI tract to perform critical life functions such as peristalsis, gut hormone secretions, and water balance during gut homeostasis (Furness, 2006). The ENS consists of various enteric neuron and glial cell types, which are classified based on molecular, electrophysiological and morphological means. In humans, along the entire ENS length, spanning from the esophagus to the anus, the ENS contains upwards of 600 million neurons, which together are capable of autonomous reflex activity (Furness, 2006), separate from the central nervous system (CNS).

In jawed vertebrates, the ENS is largely derived from neural crest cells (NCCs), a migratory and multipotent stem cell population that arises along the dorsal neural tube during neurulation within the vertebrate embryo. Specifically, the majority of the ENS is derived from NCCs that originate from a post-otic anatomical area along the neuraxis known as the vagal region (Harris and Erickson, 2007; Hutchins et al., 2018; Kuo and Erickson, 2011; Le Douarin and Teillet, 1973; Shepherd and Eisen, 2011). In zebrafish, vagal NCCs emigrate from the vagal region into anterior foregut mesenchyme tissue and migrate posteriorly down the gut length between 36 and 72 hours post fertilization (hpf) (Ganz, 2018; Harrison et al., 2014; Nikaido et al., 2018; Shepherd and Eisen, 2011; Uribe and Bronner, 2015). In all jawed vertebrates, once resident in the gut the NCCs are known as enteric neural crest cells (ENCCs) and are characterized by combinatorial expression of the genes *Sox10, Phox2b, Ret*, and *Gfra1* (Harrison et al., 2014; Hockley et al., 2019; Howard, Baker et al., 2021; Nagy and Goldstein, 2017; Shepherd et al., 2004; Southard-Smith et al., 1998; Taylor et al., 2016; Theveneau and Mayor, 2011). Following their gut colonization, zebrafish ENCCs differentiate into enteric neurons as early as 54 hpf, continuing between 72 and 120 hpf along the foregut, midgut and hindgut (Olden et al., 2008; Olsson et al., 2008). While the general phases of ENS development and gut colonization have been mapped out, to date we lack comprehensive understanding of the cellular and intercellular mechanisms that orchestrate early ENS pattern manifestation.

Several critical steps are known to be required for ENS development along the gut— including migration, proliferation and differentiation of ENCCs. Failure of ENCCs to colonize and differentiate into ENS leads to the congenital condition Hirschsprung disease (HSCR), in which variable regions of the gut lack neurons and glia in humans (Amiel et al., 2008; Heanue and Pachnis, 2007). Previous research focusing on the factors that may guide ENCC migration and their gut colonization have made recent progress. For instance, different classes of receptor tyrosine kinases (RTKs), and their corresponding ligands, have been identified as crucial components. Among these, the ENCC co-receptor Ret and its mesenchymal-derived ligand, Glial-derived neurotrophic factor (GDNF), are necessary for proper ENCC migration and survival along the gut (Landman et al., 2007; Natarajan et al., 2002; Shepherd et al., 2004; Taraviras, 1999; Young et al., 2001). Mutations in *Ret* are the leading known cause of HSCR in humans (Amiel et al., 2008). Accordingly, due to its highly prevalent involvement in HSCR manifestation, *Ret* has been the primary gene of interest for many studies, mutations in which have previously been shown to induce HSCR-like hypoganglionosis and aganglionosis, in both zebrafish and mice models (Heanue et al., 2016; Uesaka et al., 2008). However, how Ret regulates ENCC mechanisms such as proliferation, differentiation, and cell type specification *in vivo* remains poorly understood due to the highly dynamic development of ENS, GI inaccessibility of traditional amniote ENS models, as well as the ethical and technical limitations of investigating human prenatal development.

Despite advancements in our understanding of how early ENS develops, we still do not fully understand the tissue-wide strategies that ENCCs utilize to sculpt the ENS. Previous work in zebrafish (Harrison et al., 2014; Kuwata et al., 2019) have contributed to a growing body of work that implicate ENCC proliferation as key components of cell migration during colonization (Barlow et al., 2008; Peters-Van Der Sanden et al., 1993; Simpson et al., 2007; Young et al., 2004). However, we still do not fully understand the temporal dynamics of ENCC proliferation and migration during the entire process of colonization, and how those mechanisms are regulated by Ret. Here, using zebrafish larvae, the goal of this study was to resolve and quantify the cellular mechanisms of ENCC migration and proliferation in the context of Ret loss of function, in order to inform our understanding of how the ENS is constructed during early development.

## Results

### Single-cell generation tracing of ENCCs during early ENS development *in toto*

To capture the entire process of gut colonization *in vivo*, time-lapse confocal microscopy was utilized to image larval zebrafish guts *in toto* and track *-8*.*3phox2bb:*Kaede^+^ cells, which faithfully labels ENCCs (Harrison et al., 2014; Howard et al., 2022). All ENCCs were tracked, including new ENCCs generated from cell division. ENCC migration was followed from the foregut to hindgut for 48 hours, between 48-96 hpf. (**Fig. 1A-C, Movie 1**). At 48 hpf, - *8*.*3phox2bb:*Kaede^+^ ENCCs emigrated into the foregut, and migrated posteriorly until reaching the hindgut boundary at 72 hpf (**Fig. 1A,B; arrowheads**). Between 72-96 hpf, ENCCs increased in number and began patterning into nascent ENS along the gut tube (**Fig. 1B,C**).

**Figure 1.**
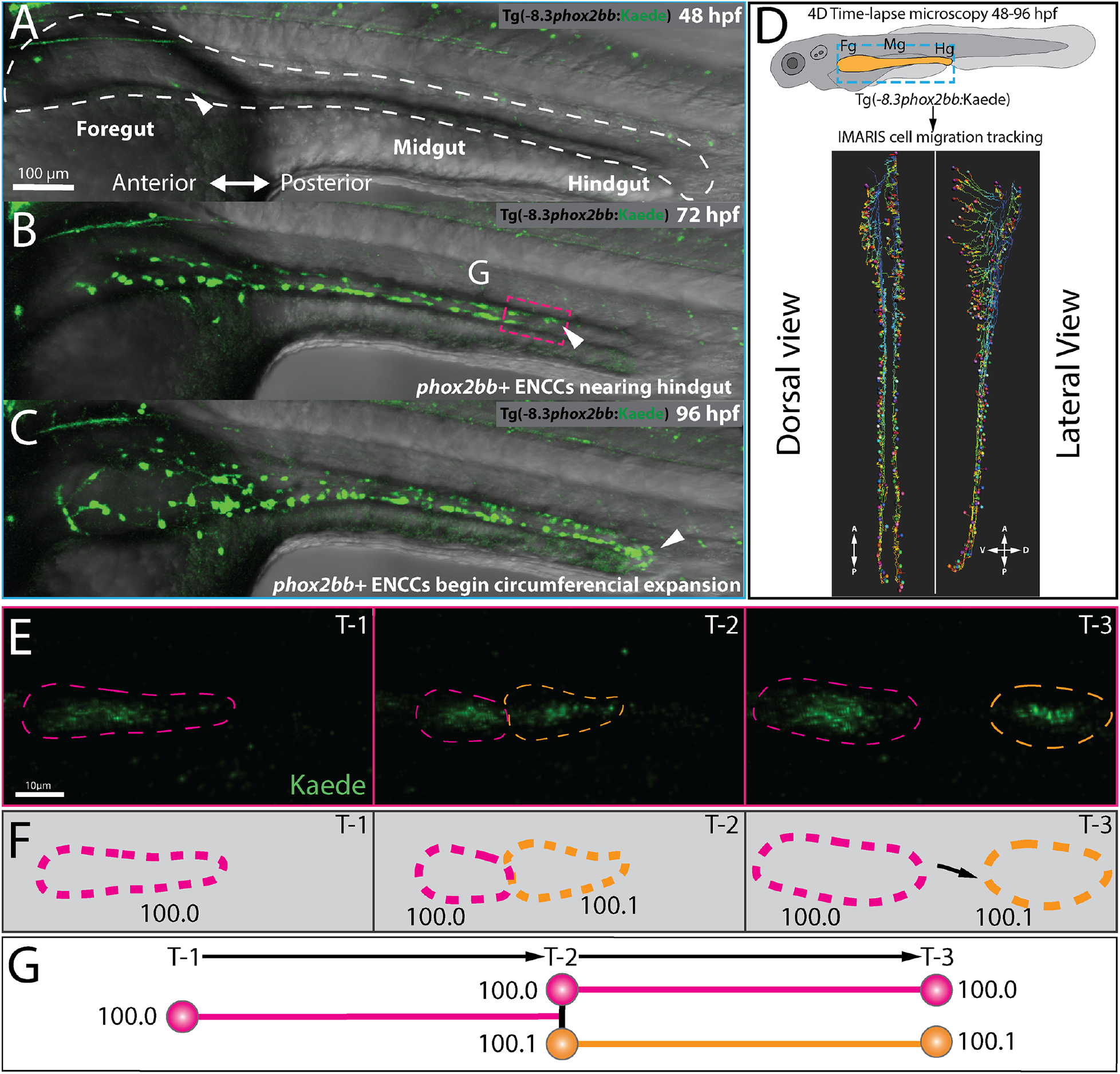
Single-cell generation tracing of ENCCs during early ENS development *in toto*. (A-C) Still images from time-lapse reveal (A, white arrow) *phox2bb*^+^ ENCCs emerging into foregut intestine at 48hpf (B, white arrow), leading ENCCs reaching the hindgut at 72 hpf (C), and the circumferential expansion of ENCCs at 96 hpf. (D) Schematic illustrates region of interests for time-lapse microscopy of ENCC migration in intestine and lateral and dorsal view snapshots of single-cell migration tracking in IMARIS. (E) Expanded images of cell migration and division at the ENCC wavefront seen in panel B, magenta box. Time-points 1-3 (T1-3) depict representative time-points where cell divisions occur. (F) Cartoon corresponds to cell outlines in panels E and with representative unique ID labels that denote cell generation number as a decimal. (G) Cartoon Generation-tree corresponds to cell divisions seen in panels E and F and depict how unique IDs are applied to cells throughout the time-lapse. A: anterior; P: posterior; D: dorsal; V: ventral, shown in D.

IMARIS image analysis software allowed us to manually track the generations of every ENCC during the 48-hour time period (**Fig. 1D, Movie 2**). Tracking including newly generated ENCCs produced by cell division as shown in spatial cell tracks along the gut tube (**Fig. 1D; Movie 3,4)**, and in an enteric generation tree over time (**Fig. S1**). IMARIS tracking produced unique cellular generation labels that denoted ENCC hierarchical order within their respective generation tree, as schematized in **Fig. 1E-G**. As a cell divided along the anterior-posterior gut axis, the anterior most daughter cell retained the unique ID and generation label of the original parent cell, while the posterior daughter cell was given a unique ID with a generation label corresponding to +1 to that of the parent cell that gave rise to the daughter (**Fig. 1F,G**). This imaging and generation analysis pipeline shows that zebrafish ENCC populations can be followed at single cell resolution over time and sets the stage to scrutinize gut-scale events during normal and abnormal early ENS development.

### A novel CRISPR-Cas9 generated *ret* zebrafish mutant

In order to study the cellular mechanisms underlying ENS formation, and inform studies aimed at understanding cellular basis for neurocristopathy manifestation, the evolutionarily conserved gene, *ret*, was targeted for CRISPR-Cas9 mutagenesis. To accomplish this, a single guide RNA (sgRNA) targeting the 8^th^ exon of zebrafish *ret* (**Fig. 2A**) was generated and injected along with *cas9* mRNA into *-8*.*3phox2bb*:Kaede embryos (**Fig. 2B**). The mutation was isolated and identified by outcrossing injected F0 *-8*.*3phox2bb*:Kaede fish to AB WT to produce F1 families (**Fig. 2B**). F1 fish were fin clipped for genomic DNA isolation that was then sequenced; this identified an 11 base pair (bp) deletion resulting in a predicted premature stop codon within the coding sequence (CDS) (**Fig. 2C**), which we designate hereafter as *ret*^*wmr1*^. Following in-cross of *ret*^*wmr1/+*^ F1s, F2 were seen to display either: WT (*ret*^*+/+*^) (**Fig. 2D**), hypoganglionosis (HSCR-like) (*ret*^*wmr1/+*^) (**Fig. 2E**), or total aganglionosis (*ret*^*wmr1/wmr1*^) (**Fig. 2F**) phenotypes. HSCR-like phenotypic fish were found in less than 50% F2 progeny due to incomplete penetrance and considerable variability in extent of aganglionosis (**Table S1**), an observation that is congruent with human HSCR manifestation, mammalian HSCR models, and other *ret* heterozygous zebrafish models (Heanue and Pachnis, 2007; Heanue et al., 2016; Lake and Heuckeroth, 2013; Stanchina et al., 2006). To assess whether *ret*^*wmr1*^ retained any of its function, a rescue experiment was performed by injection of mRNA coding for *ret*^*wmr1*^, *or ret*^*WT*^, into F2 incrossed *-8*.*3phox2bb*:Kaede;*ret*^*wmr1/+*^ embryos (**Fig. S2**). Injection of *ret*^*wmr1*^ mRNA failed to rescue the number of larvae exhibiting total aganglionosis phenotypes by 96 hpf while *ret*^*WT*^ mRNA partially rescued diseased phenotypes, suggesting functional loss of *ret*^*wmr1*^ mutant. The successful generation of the *ret*^*wmr1/+*^ fishline provides a valuable zebrafish model for studying fundamental ENCC dynamics and ENS development.

**Figure 2.**
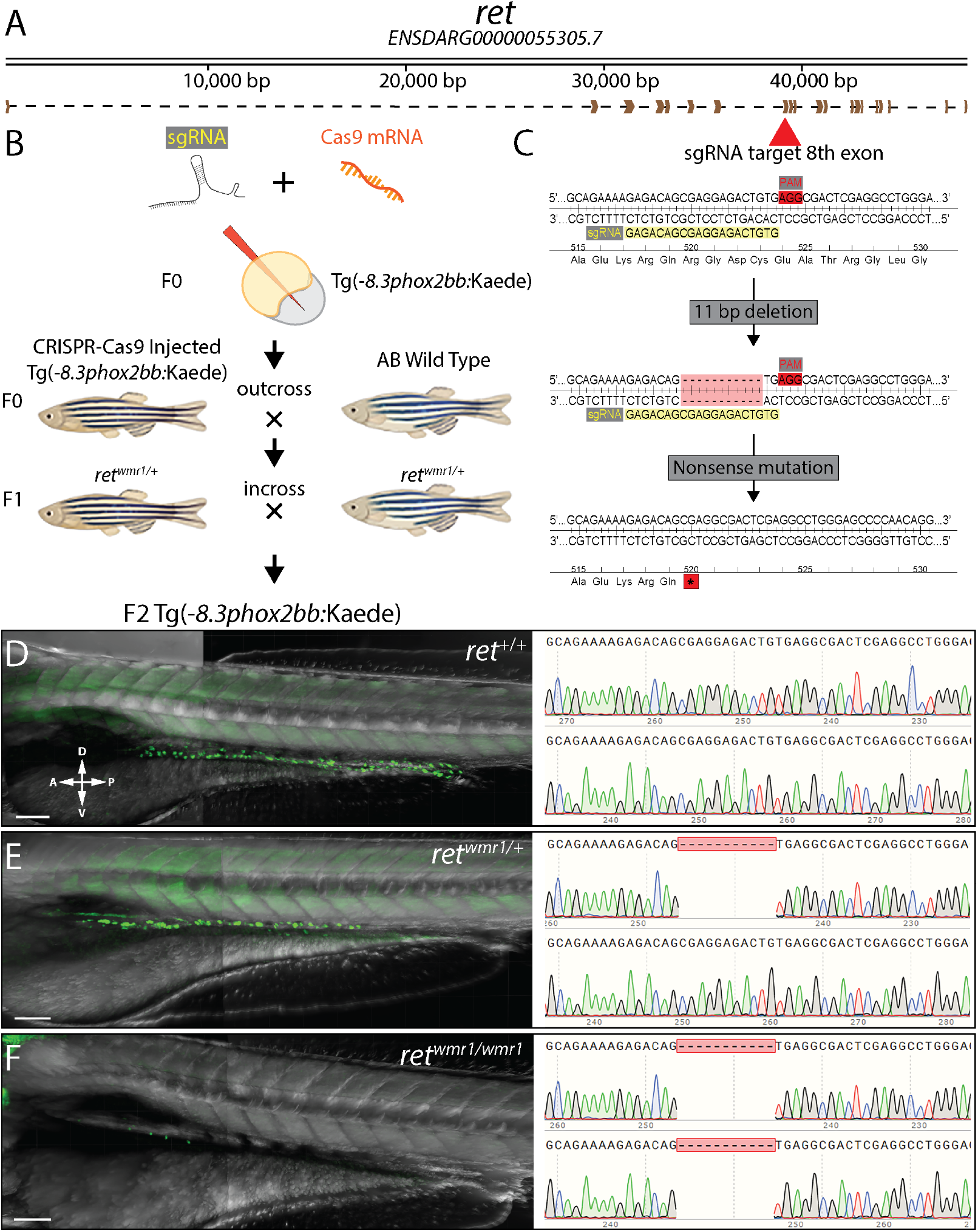
A novel CRISPR-Cas9 generated *ret* zebrafish mutant. (A) Schematic of receptor tyrosine kinase gene, *ret* and its resulting mRNA. (B) Diagram depicting injection of sgRNA and Cas9 mRNA into *ret*^*+/+*^ Tg(−8.3*phox2bb*:Kaede) and proceeding crosses used to isolate single CRISPR generate lesion within *ret* allele. (C) Zoomed in region of coding sequence of 8^th^ exon (red arrow) targeted by sgRNA sequence (GAGACAGCGAGGAGACTGTG, yellow highlight) adjacent to pam motif (red highlight). CRISPR-Cas9 activity at this locus generated an 11bp deletion that resulted in nonsense mutation (Stop codon, red asterisk). (D-F) Resulting genotypes from in-crossing F1 heterozygotes harboring nonsense mutation (*ret*^*wmr1/+*^) produces phenotypic (D) WT (*ret*^*+/+*^) (E), Hypoganglionosis or HSCR-like (*ret*^*wmr1/+*^) and (F), total aganglionosis (*ret*^*wmr1/ wmr1*^). Scale bars = 100 *µ*m. A: anterior; P: posterior; D: dorsal; V: ventral, shown in D.

### *ret*^*wmr1/+*^ fish display a reduction in total ENCC number and migratory extent along the gut

To examine how the enteric defect phenotypes manifest in the *ret*^*wmr1/+*^ model, we utilized the *in toto* time-lapse microscopy paradigm, as described in Figure 1. Time-lapse datasets of *ret*^+/+^ (**Fig. 3A, Movie 1**) and *ret*^*wmr1/+*^ (**Fig. 3B, Movie 5**) fish were used to quantify total ENCC numbers over time, between 48-96 hpf (**Fig. 3C**). While both conditions showed a steady temporal increase in ENCC number, *ret*^*wmr1/+*^ fish displayed a significant reduction in overall ENCC number; while controls displayed an average of 172 ENCCs by 96 hpf, heterozygotes only had 101 (**Fig. 3C**). In each condition, ENCCs were observed migrating in two parallel chains (left and right) on lateral sides of the gut tube (**Fig. 3D,E,H,I; Movie 2 and 6**). The distance traveled by the leading most ENCCs (vanguard cells), on the left and right migratory chain, was averaged across replicates and mapped in relation to the foregut region throughout the time course, for both *ret*^*+/+*^ and *ret*^*wmr1/+*^ conditions (**Fig. 3F**). In both conditions, vanguard cells in the left and right migratory chains progressed posteriorly at similar rates until they reached their final distance at 96 hpf (**Fig. 3F,G**). *ret*^*+/+*^ vanguard ENCCs displaced in a near linear fashion over time until they reached the distal boundary of the gut tube at ∼78 hpf, approximately 1000 *µ*m from the foregut, at which point they could migrate no further (**Fig. 3D, F-H**). In contrast, while *ret*^*wmr1/+*^ ENCCs migrated down the gut length, they failed to reach the hindgut and migrated only ∼650 *µ*m from the foregut, a significantly shorter migration distance than *ret*^*+/+*^ vanguard cells (**Fig. 3E,F,G,I**). Collectively, these data demonstrate a reduction in overall ENCC number and migratory extent between 48-96 hpf in *ret*^*wmr1/+*^ fish. These results suggest that the collective cell migration and displacement of leading edge ENCCs may be directly correlated to overall ENCC number.

**Figure 3.**
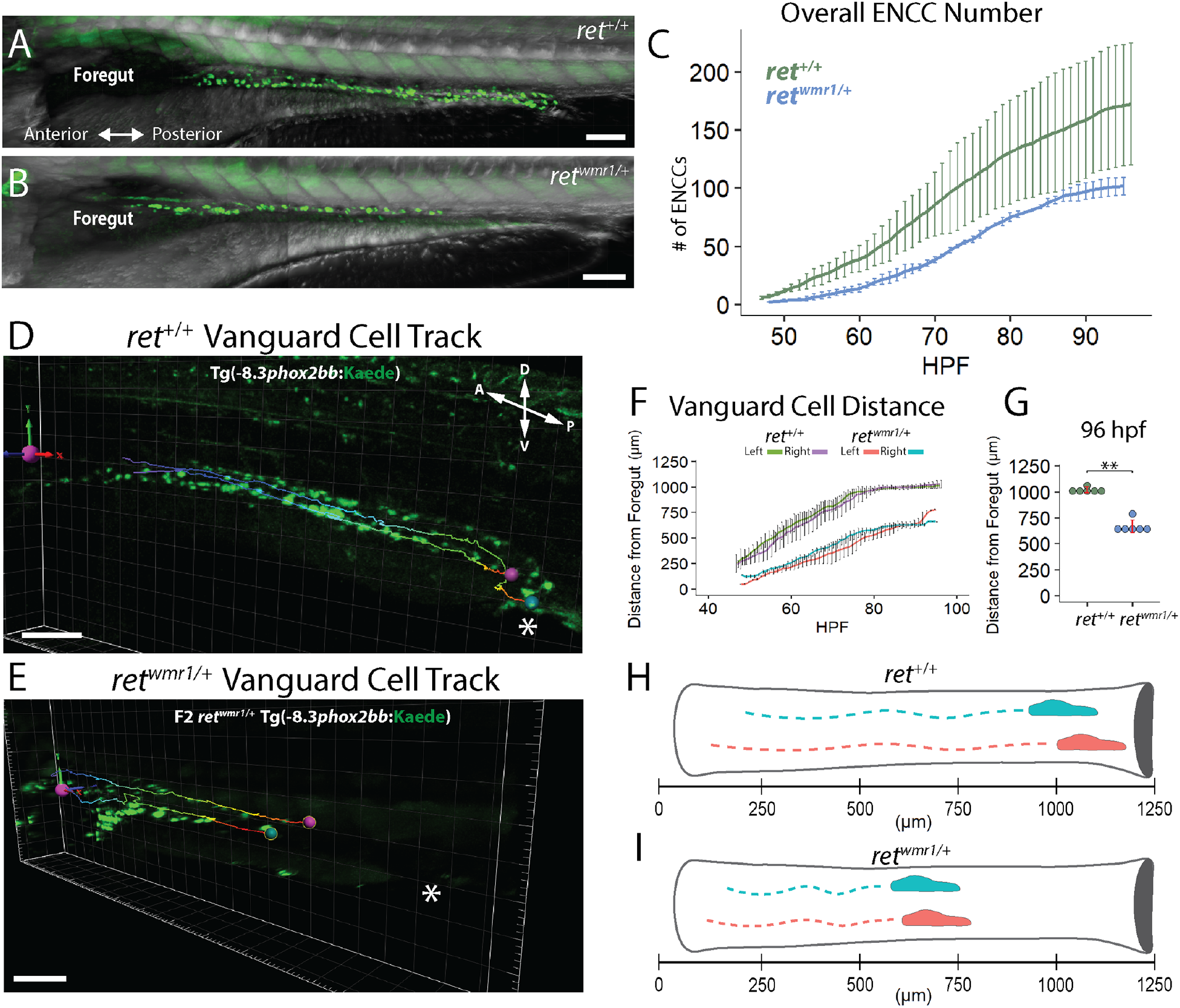
*ret*^*wmr1/+*^ fish display a reduction in total ENCC number and migratory extent along the gut. (A and B) Image of (A) *ret*^*+/+*^ and (B) *ret*^*wmr1/+*^ Tg(−8.3*phox2bb*:Kaede) fish show *phox2bb*^+^ ENCCs within the intestine at 96 hpf. *ret*^*wmr1/+*^ ENCCs fail to reach distal hindgut at 96 hpf. (C) Average number of ENCCs present during the 48-96 hpf time-lapse in *ret*^*+/+*^ and *ret*^*wmr1/+*^ fish (N=3). (D and E) Snapshot of final 96 hpf time-point depicts migratory track of leading-edge cell (Vanguard) in left (pink spot) and right (blue spot) ENCC chains (Cloaca=white asterisk). (F) Average distance traveled by left and right Vanguard cells in *ret*^*+/+*^ and *ret*^*wmr1/+*^ fish compared across 48-96 hpf (N = 6 per condition). (G) Final position of Vanguard Cells at 96 hpf (Wilcox Test, *P =* 0.0022). (H and I) Cartoon depicts Vanguard cells in left (pink) and right (blue) ENCC migratory chains in (H) *ret*^*+/+*^ and (I) *ret*^*wmr1/+*^. Scale bars = 100 *µ*m. A: anterior; P: posterior; D: dorsal; V: ventral, shown in D.

### Generation of new ENCCs along the gut is compromised in *ret*^*wmr1/+*^ fish

To investigate the hypothesis that ENCC number is correlated to cell displacement, we aimed to determine how the production of ENCC generations via cell proliferation was related to the overall displacement and spatiotemporal patterning of cells along the gut length. Using previously described generation labels (**Fig. 1F,G**), the total number of ENCC generations, as well as distances from the foregut of individual ENCC generations, were averaged and plotted versus hpf in *ret*^*+/+*^ (**Fig. 4A**) and *ret*^*wmr1/+*^ (**Fig. 4C**) conditions. In *ret*^*+/+*^ fish, ENCCs produced 18 generations between 48-96 hpf, where each subsequent generation migrated further posterior than their predecessors in an ordered fashion until the gut tube was colonized (**Fig. 4A,B**). While *ret*^*wmr1/+*^ ENCCs displayed similar posterior patterning, with subsequent generations migrating further than their predecessors, these ENCCs produced only 13 generations that failed to colonize the full gut length (**Fig. 4C,D**). Plotting the average number of cells per given generation revealed that *ret*^*wmr1/+*^ ENCC generations contained fewer cells, when compared to *ret*^*+/+*^ (**Fig. 4E**). These data support a model of proliferation driven migration, where continual cell division produces daughter cells that migrate further than their predecessors, and suggests that reducing the number of cell divisions reduces their migratory extent.

**Figure 4.**
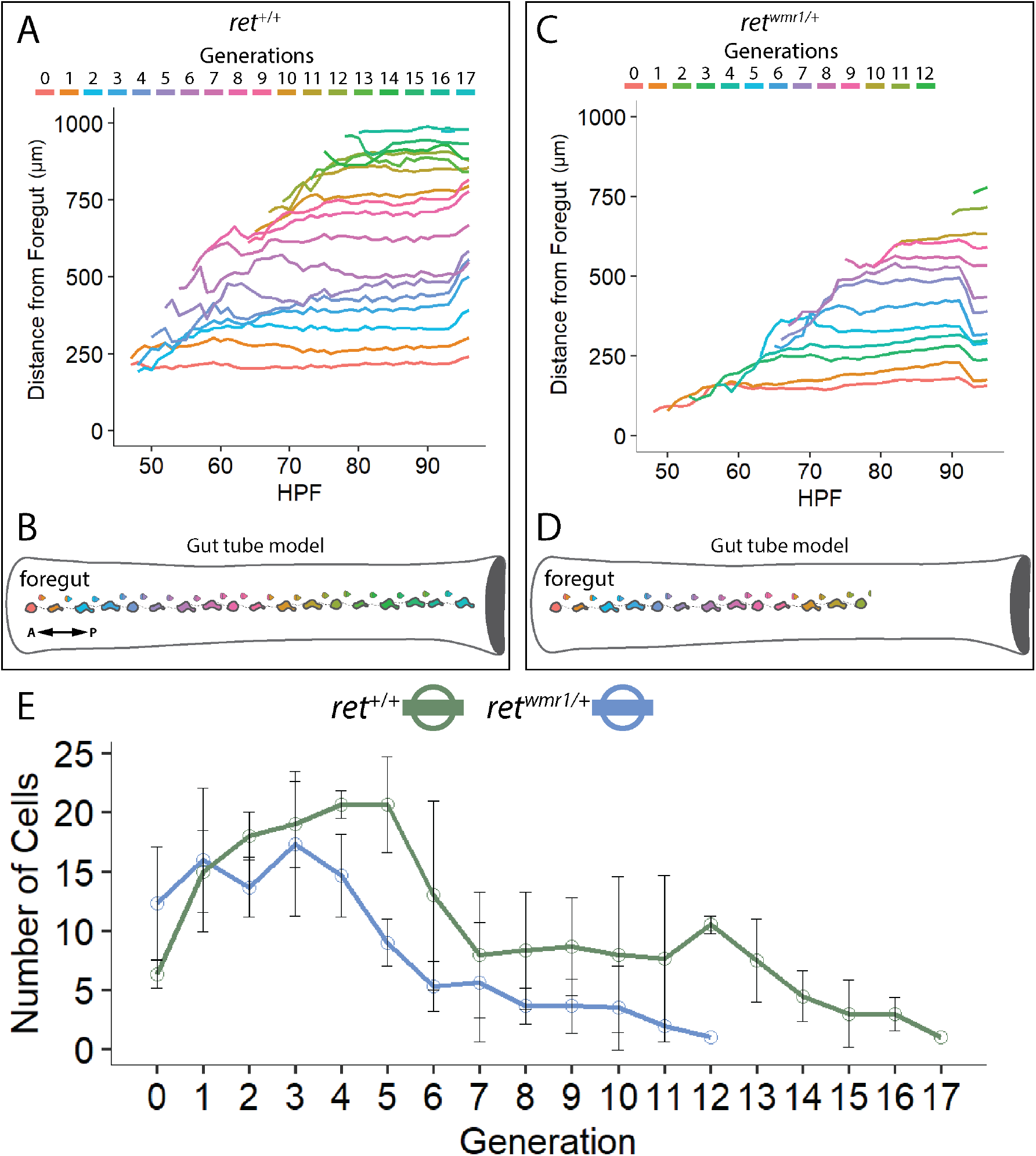
Generation of new ENCCs along the gut is compromised in *ret*^*wmr1/+*^ fish. (A-D) ENCC proliferation produces successive generations of cells that migrate further than predecessors along the anterior-posterior axis of the gut tube. (A and B) *ret*^*+/+*^ ENCCs produce up to 18 generations to effectively colonize the entire gut tube (N=3). (C and D) *ret*^*wmr1/+*^ ENCCs produce up to 13 generations that successively colonize the gut tube but fail to reach the hindgut (N=3). (E) Average number of cells per given generation are reduced in *ret*^*wmr1/+*^ fish when compared to *ret*^*+/+*^. A: anterior; P: posterior, shown in B.

### Analysis of *ret*^*wmr1/+*^ highlights that distance between ENCCs is associated with deficient migration and proliferation during ENS formation

Based on our observation that cell division was associated with cell displacement overtime, we expected to see a reduction in cell migration speed as a function of cell number during this same window of ENCC migration. To quantify advancement rates of the leading most ENCCs between 48-96 hpf, the migratory speed of vanguard ENCCs were averaged across conditional replicates (**Fig. 5A, B**). A sliding mean curve (every 5 hours) was used to visualize average vanguard speed in *ret*^*+/+*^ (green) and *ret*^*wmr1/+*^ (blue) conditions (**Fig. 5A**). While ENCCs in both conditions displayed stable speeds between 48-70 hpf, *ret*^*wmr1/+*^ ENCCs showed lower speeds (∼21 *µ*m/hr), when compared to *ret*^*+/+*^ (∼34 *µ*m/hr) (**Fig. 5A**). Indeed, box plots comparing the average vanguard speed at intervals between 48 and 96 hpf revealed that *ret*^*wmr1/+*^ ENCCs migrated significantly slower than *ret*^*+/+*^ specifically during the 48-60 hpf interval (**Fig. 5B**).

**Figure 5.**
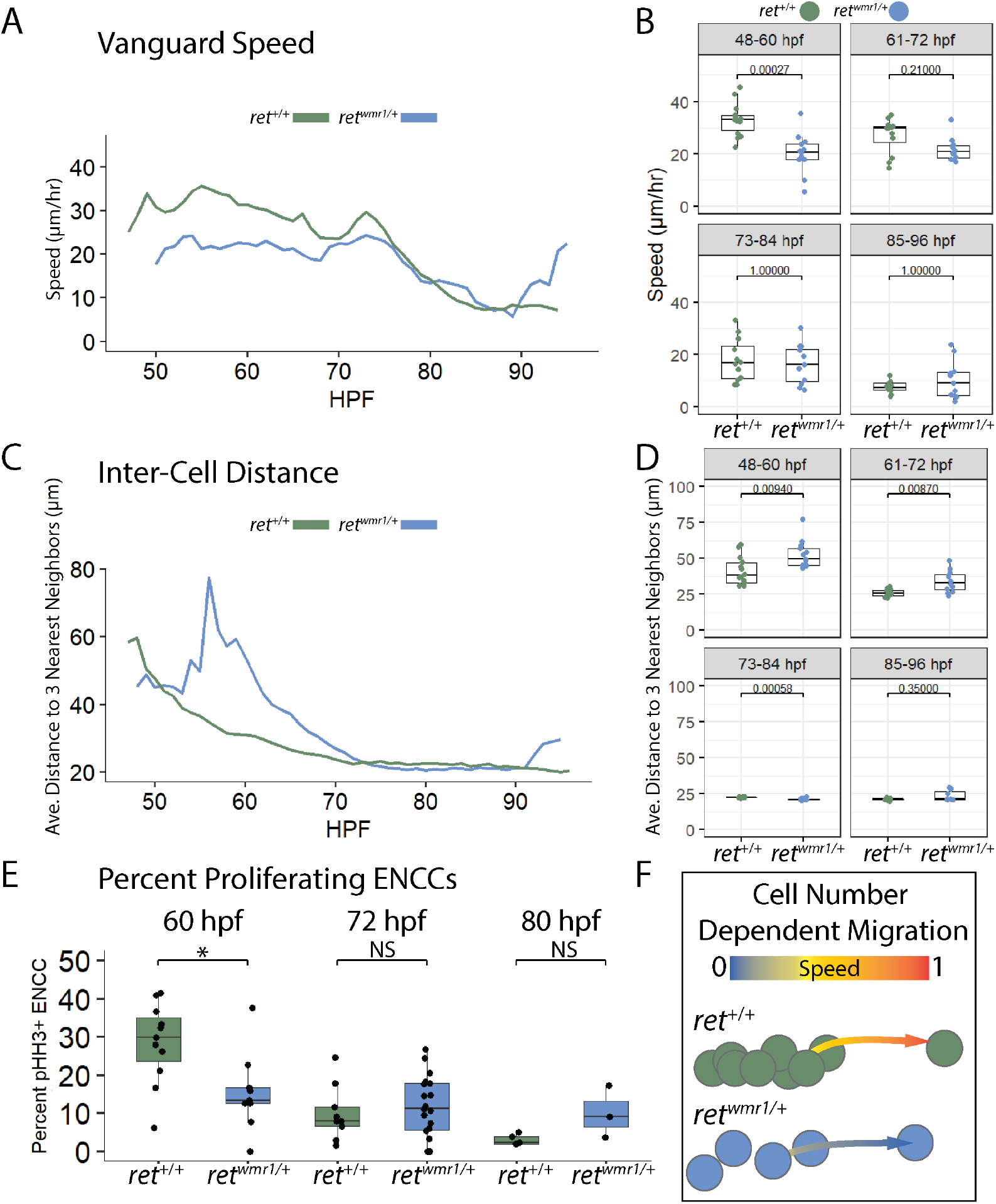
Proliferation deficient *ret*^*wmr1/+*^ ENCCs highlight cell density associated migration mechanism during ENS formation. (A and B) Average speed of vanguard cell migration reveals a reduction in *ret*^*wmr1/+*^ ENCC speed during key cell migration window 48-60 hpf (N=6 per condition; Wilcox.test, *P =* 0.00027). (C and D) Average Distance to 3 nearest neighbors was calculated for each ENCC within an individual population and then averaged across conditional replicates and plotted vs hpf for both *ret*^*+/+*^ and *ret*^*wmr1/+*^ fish (N = 3 per condition). Significantly higher distances between neighbors were observed among *ret*^*wmr1/+*^ ENCCs between 48-60 hpf (Wilcox.test, *P =* 0.00940) and 61-72 hpf (Wilcox.test, *P* = 0.00870), indicative of lower cell density. (E) Percent proliferating ENCCs was calculated using whole-mount immunofluorescence performed using Tg(−8.3*phox2bb*:Kaede) fish in combination with anti-pHH3 and anti-Kaede antibodies to calculate the percent pHH3^+^, Kaede^+^ ENCCs at 60 hpf, 72 hpf, and 80 hpf. *ret*^*wmr1/+*^ fish displayed lower percent of proliferating ENCCs at 60 hpf (Wilcox. Test, *P =* 0.02). (F) Graphic depicts density dependent migration model where more densely spaced *ret*^*+/+*^ ENCCs display higher speeds of migration than less densely spaced *ret*^*wmr1/+*^ ENCCs.

In addition to slower speeds, in time-lapse movies we noted that ENCCs localized along the gut tube of *ret*^*wmr1/+*^ mutants looked qualitatively less densely spaced than *ret*^*+/+*^ control fish. To measure distances between ENCC along the gut during their migration, the average distance between three nearest ENCC neighbors was calculated and averaged across conditional replicates (**Fig. 5C**). While the *ret*^*wmr1/+*^ ENCCs eventually reached a similar average inter-cell distance as *ret*^*+/+*^ ENCCs by ∼72 hpf (∼24 *µ*m), *ret*^*wmr1/+*^ ENCCs were found to have greater inter-cell distances between ∼48-72 hpf (∼49 *µ*m vs ∼38 *µ*m), congruent with the migratory window in which *ret*^*wmr1/+*^ ENCCs displayed slower migration speeds (**Fig. 5B**). Confirming this observation, box plots comparing the average distance between three nearest neighbors during windows at 48-60 hpf and 61-72 hpf showed that *ret*^*wmr1/+*^ ENCCs were spaced significantly further apart than their *ret*^*+/+*^ counterparts (**Fig. 5D**).

To further investigate the mechanism underlying lower vanguard displacement speed and overall ENCC inter-cell distances, we next sought to quantify ENCC proliferation *in situ*. Towards this end, a phospho-histone H3 (pHH3) antibody was used on fixed tissue at key time points of 60, 72, and 80 hpf to determine the percent of ENCCs that were actively proliferating (**Fig. 5E**) (Gurley et al., 1978; Hendzel et al., 1997; Pérez-Cadahía et al., 2009). From these data, a lower percentage of ENCCs were pHH3^+^ in *ret*^*wmr1/+*^ fish at 60 hpf, when compared with *ret*^*+/+*^ ENCCs, while there was no significant difference at 72 and 80 hpf (**Fig. 5E**).

Collectively, these above-described data regarding cell speed, inter-cell distances and proliferation highlight an important developmental window between 48-72 hpf, in which *ret*^*wmr1/+*^ ENCCs exhibit their most significant deficiencies. These results suggest a model of proliferation coupled migration, and/or inter-cell distance dependent migration, where proliferation deficient ENCCs fail to reach a cell number threshold needed to sustain posterior displacement of ENCCs along the length of the gut (**Fig. 5F**).

### *ret*^*wmr1/+*^ ENCCs display accelerated neuron differentiation gene expression and immunoreactive signatures

Single-cell sequencing data from our prior work revealed that ENCCs displayed lineage-specific neuronal gene expression patterns at 69 hpf (Howard, Baker et al., 2021). Specifically, in addition to pan neuronal markers, such as *elavl3*, ENCCs were beginning to differentially express unique combinations of neurochemical gene markers; such as, *vipb, nos1*, and *slc18a3a*, as well as the Intrinsic Primary Afferent Neuron (IPAN) specific transcription factor, *pbx3b* (Howard, Baker et al., 2021). Notably, *vipb* was found to be expressed in nearly all early differentiating ENCCs, while the IPAN marker *pbx3b* was found in a very small subset of ENCCs, denoting a more specific enteric neuron subtype that arises later during ENS development. Due to the spatiotemporal specific emergence of these enteric markers, we hypothesized that the timing and levels of their expression would be altered within *ret*^*wmr1/+*^ fish, indicative of aberrant differentiation timing.

In order to investigate the above hypothesis, hybridization chain reaction (HCR) was used to investigate gene expression patterns of *ret*^*wmr1/+*^ ENCCs, in comparison to *ret*^*+/+*^, at the developmental time-points between 60-120 hpf (**Fig. 6A-I**). Due to its near ubiquitous expression, probes against *vipb* transcript were used to label differentiating enteric neurons, while *pbx3b* was used to mark enteric neuron specification into differentiating IPANs. From these analyses, *ret*^*wmr1/+*^ fish exhibited a higher percent of ENCCs displaying an IPAN gene signature, when compared with *ret*^*+/+*^ cells, with the peak discrepancy appearing at 96 hpf (**Fig. 6E-G**). However, by 120 hpf *ret*^*wmr1/+*^ and *ret*^*+/+*^ fish exhibited roughly an equal percentage (∼40%) of ENCCs with IPAN signature within the total population of *vipb*^+^ enteric neurons, suggesting premature enteric neuron specification rather than altered subtype-specification (**Fig. 6H-J**). To determine if *pbx3b* transcript levels were differentially expressed between the conditions, total mean *pbx3b* pixel intensity was calculated throughout the entire gut tube tissue to approximate its total expression within the ENS, irrespective of its co-expression with *vipb* (**Fig. 6K**). Corroborating our prior *pbx3b*^+^ and *vipb*^+^ ENCC cell counts, the data demonstrated that *ret*^*wmr1/+*^ fish exhibited significantly higher levels of *pbx3b* expression than *ret*^*+/+*^ tissue as early as 72 hpf, with the greatest discrepancy occurring at 96 hpf (**Fig. 6K**). Moreover, while the fraction of *pbx3b*^+^ ENCCs was higher in *ret*^*wmr1/+*^ fish at 96 hpf, no correlation was found between the fraction of *pbx3b*^+^ ENCCs and the extent of aganglionosis in HSCR-like larvae (**Fig. S3)**. These findings suggest Ret signaling serves to regulate the timing in which ENCCs progress through their differentiation program, and consequentially their proliferative potential.

**Figure 6.**
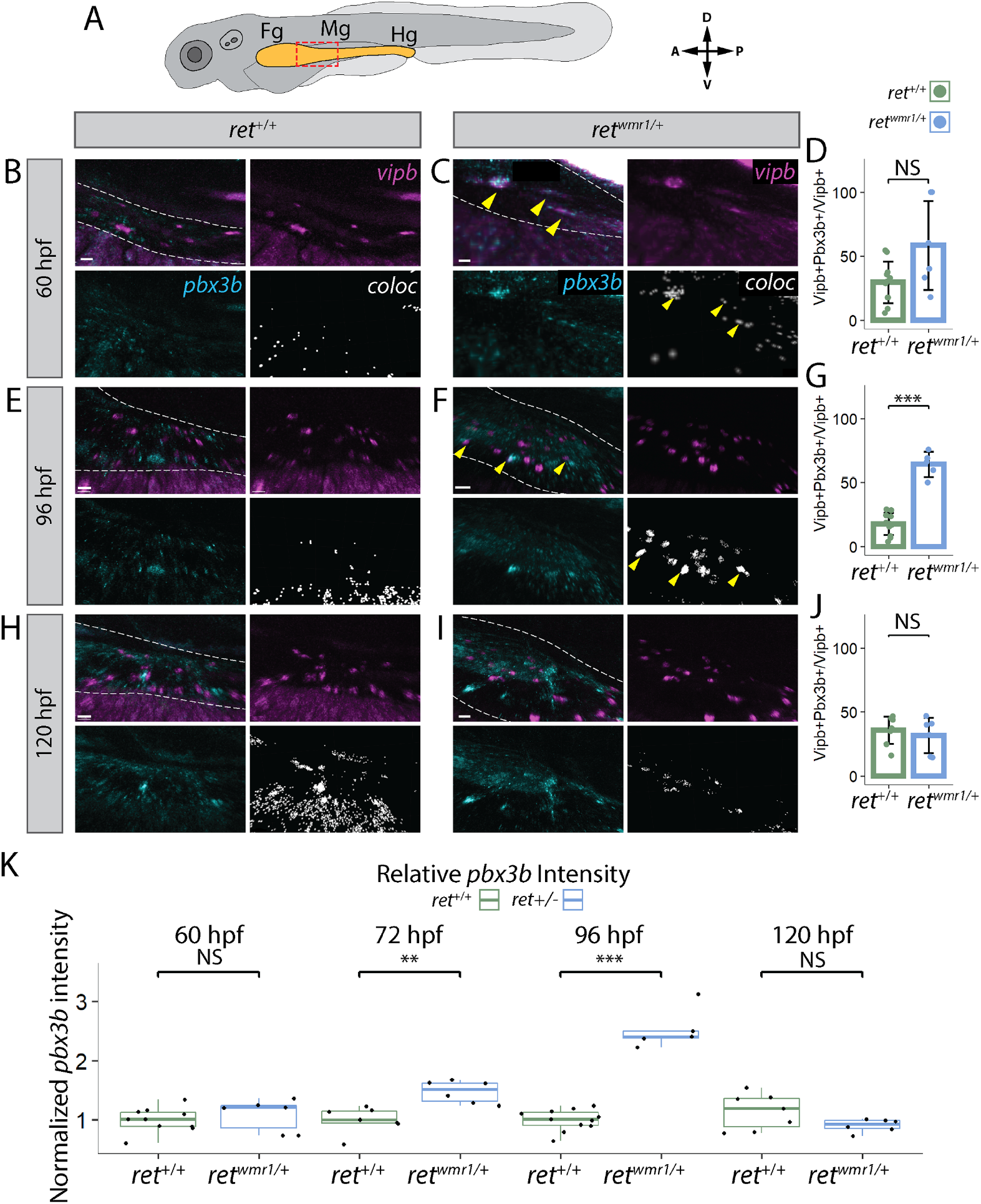
*ret*^*wmr1/+*^ ENCCs display accelerated neuron differentiation gene expression. (A-K) Hybridization Chain Reaction (HCR) performed using probes targeting *pbx3b* and *vipb* transcripts. (A) Cartoon larvae denotes foregut (Fg), midgut (Mg), and hindgut (Hg) intestine (yellow) with red box indicating foregut-midgut boundary region of interest from which representative images are displayed. Representative images reveal *vipb* (magenta), *pbx3b* (cyan) expression, and their colocalized (coloc) channel (white) within gut tube (white dashed outline) at (B and C) 60 hpf, (E and F) 96 hpf, and (H and I) 120 hpf. Yellow arrows indicate co-positive *vipb*^*+*^*/pbx3b*^*+*^ ENCCs. (D,G, and J) Percent co-positive *vipb*^*+*^*/pbx3b*^*+*^ calculated from total *vipb*^*+*^ ENCCs at (D) 60 hpf, (G) 96 hpf (Wilcox.test, *P =* 0.00046), and (J) 120 hpf. (K) Mean *pbx3b* channel intensity quantified throughout entire gut tube and normalized to *ret*^*+/+*^ average intensity at 60 hpf (Wilcox.test, NS), 72 hpf (Wilcox.test, *P =* 0.00117), 96hpf (Wilcox.test, *P =* 0.00046), and 120 hpf (Wilcox.test, NS). (B, C and I) Scale bar = 20 *µ*m. (E) Scale bar = 15 *µ*m. (F) Scale bar = 30 *µ*m. (H) Scale bar = 10 *µ*m. A: anterior; P: posterior; D: dorsal; V: ventral, shown in A. In (D) *ret*^*+/+*^ N = 11, *ret*^*wmr1/+*^ N = 6, in (G) *ret*^*+/+*^ N = 11, *ret*^*wmr1/+*^ N = 5, and in (J) *ret*^*+/+*^ N = 8, *ret*^*wmr1/+*^ N = 6.

To determine if enteric neuron differentiation timing was indeed altered, as suggested by the HCR *in situ* gene expression data, whole-mount immunohistochemistry was performed (**Fig. 7A-F**) using antibodies targeting the pan-neuronal marker, Elavl3 to label all enteric neurons and, choline acetyltransferase (ChAT) to label cholinergic enteric neurons, including IPANs (Furness et al., 2004; Morarach et al., 2021). Experiments were performed on *ret*^*+/+*^ and *ret*^*wmr1/+*^ fish at the developmental time-points 96 hpf 120 hpf to compare fractions of ChAT^+^ neuron protein reactivity among Elavl3+ enteric neurons (**Fig. 7D, G**). Comparable to *pbx3b* expression fractions (Fig. 6), the percentage of ChAT^+^ cells were found to be significantly higher in *ret*^*wmr1/+*^ larvae, when compared to *ret*^*+/+*^ larvae at 96 hpf (**Fig. 7D**), while they were found to be at near equal levels by 120 hpf (**Fig. 7G**). Together, these data provide supporting evidence that a reduction in *ret* function results in accelerated enteric neuron differentiation.

**Figure 7.**
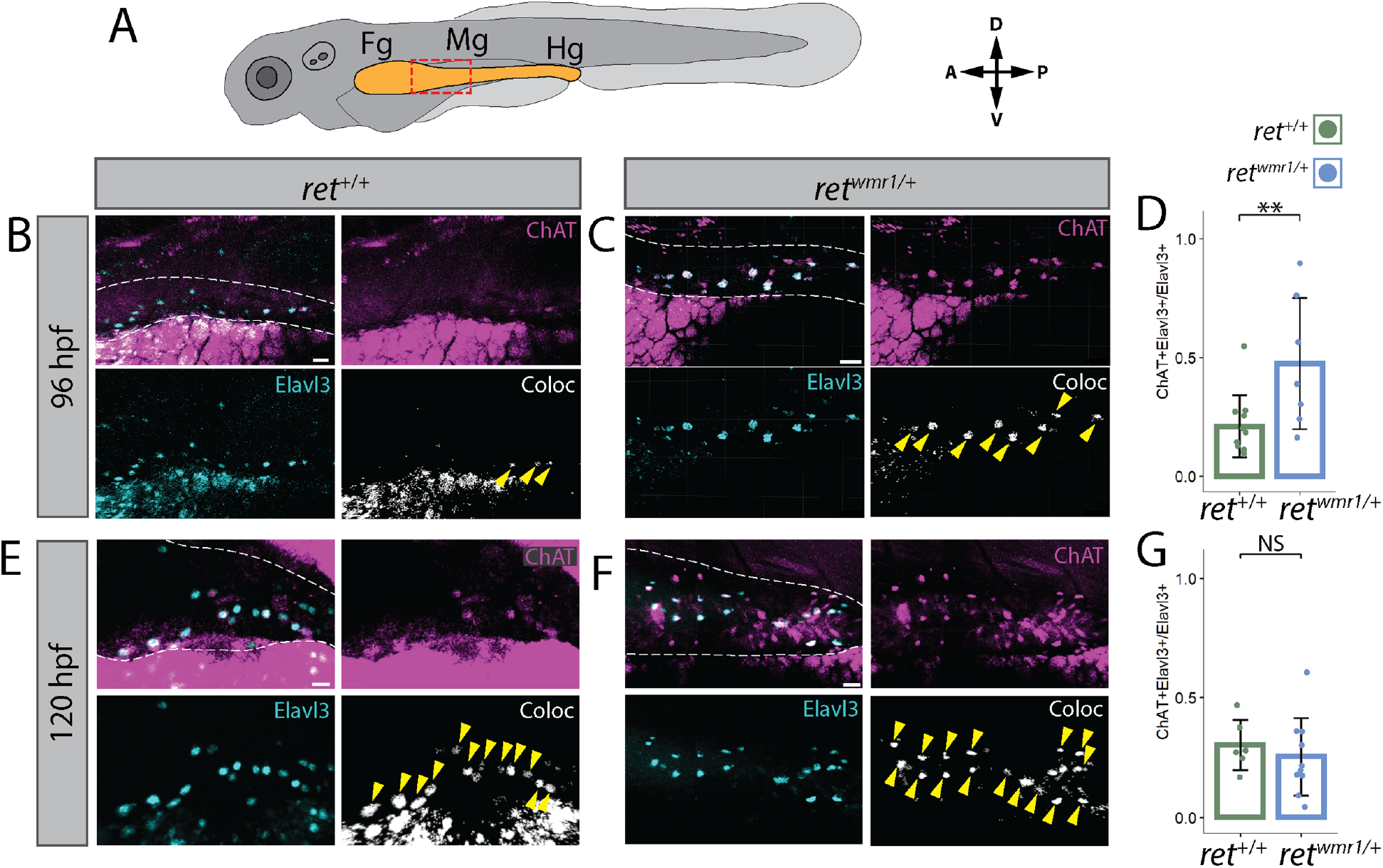
*ret*^*wmr1/+*^ ENCCs display accelerated neuron immunoreactive signatures. (A) Cartoon larvae denotes foregut (Fg), midgut (Mg), and hindgut (Hg) intestine (yellow) with red box indicating foregut-midgut boundary region of interest from which representative images are displayed. (B,C,E,F) Whole-mount immunohistochemistry performed using antibodies targeting ChAT and Elavl3. Representative images reveal ChAT (magenta), Elavl3 (cyan) expression, and their colocalized (coloc) channel (white) within gut tube (white dashed outline) at (B and C) 96 hpf, (E and F) and 120 hpf. Yellow arrows indicate co-positive ChAT^+^*/*Elavl3^+^ ENCCs. Percent co-positive ChAT^+^*/*Elavl3^+^ calculated from total Elavl3^+^ enteric neurons at (D) hpf (p=0.0083), and (G) 120 hpf (NS). (B, C and F) Scale bar = 20 *µ*m. (E) Scale bar = 15 *µ*m. In (D) *ret*^*+/+*^ N = 11, *ret*^*wmr1/+*^ N = 7, in (G) *ret*^*+/+*^ N = 6, *ret*^*wmr1/+*^ N = 10.

## Discussion

We have utilized a zebrafish model to show, for the first time, *in toto* colonization of the gut by ENCC at single-cell resolution in a vertebrate. We found over the course of 48 hours of development that several cellular and intercellular events are critical during colonization; including, ENCC number maintenance, migratory speed, ENCC spacing along the gut, and proliferative expansion. Specifically, we mapped out the generations of all ENCC cells during their migration and found that proliferative expansion is dampened during precise temporal phases along the gut in *ret*^*wmr1/+*^ mutants, greatly stalling enteric colonization and leading to severe colonic aganglionosis. Moreover, we discovered premature neuronal differentiation signatures in the context of aberrant *ret* heterozygous background, suggesting that Ret is required for proper timing of neurogenesis of ENCCs during their migration.

Previous studies performed in amniote models have demonstrated the necessity of proper ENCC numbers for gut colonization (Barlow et al., 2008; Peters-Van Der Sanden et al., 1993; Young et al., 2004). For example, in the avian ENS, manual ablation of NCCs at E1.5, prior to their entrance into gut, was sufficient to reduce the extent of ENCC colonization within the gut, highlighting the importance of proper cell numbers during the early stages of NCC migration (Barlow et al., 2008). Collectively, these studies provide supporting evidence for “population pressure” driven migration, whereby sufficient ENCC numbers are required for sustained migration and subsequent ENS development. Our present work in the anamniote zebrafish corroborates and extends these prior findings in amniotes by demonstrating the association between total ENCC number and displacement of vanguard cells along the gut completely *in toto*—providing evidence of a conserved mechanism of ENCC migration present among vertebrates. While the findings of our study appear congruent with studies performed in amniote models, it should be noted that the zebrafish ENS model has limitations when making comparisons to mammalian ENS development. Notably, the contribution of sacral-NCCs to the ENS, which has been extensively shown during mammalian ENS development (Rao and Gershon, 2018), has not been shown in zebrafish to date. Additionally, the zebrafish gut anatomy is less complex, lacking a submucosal ENS plexus, which is a crucial component of the mammalian ENS (Wallace et al., 2005).

Our data revealed a developmental window of time during which a critical number of ENCCs are required during gut colonization. Specifically, at 60 hpf when ENCCs were approximately halfway through the zebrafish gut tube (midgut), when compared with wildtype, we saw a significant reduction in ENCC proliferation in *ret*^*wmr1/+*^ mutants, the timing of which coincided with significantly increased inter-cell distances and reduced cell migration speeds. A complementary study performed in E12.5 mouse explants, a stage later in ENS development and comparable to 72 hpf in zebrafish, found that following ablation of anterior localized ENCCs, isolated vanguard ENCCs posterior to the lesion site migrated at slower speeds, leading to the conclusion that cell number and cell-cell contact were required for proper migratory wavefront speed (Young et al., 2004). Very recently, a study utilizing fixed mouse embryos found that colonization was drastically reduced/delayed in a *Ret* homozygous background, and upon examination of proliferation, discovered that ENCCs displayed reduced proliferation at E10.5 (Natarajan et al., 2022), which is roughly equivalent to zebrafish 54-60 hpf and in general alignment with our current findings. Our work at present also corroborates the conclusions of Young et al. by demonstrating a lower cell count correlated with shorter migration displacement and slower speeds. It is possible, in addition to the gut-length observation of decreased proliferation and speeds of ENCCs we saw following loss of *ret*, that early reduced ENCC numbers prior to gut entry could contribute to the observed phenotypes, which could be tested in a future study.

A recent study performed using a *ret*^*+/-*^ zebrafish model, that also presents an ENS HSCR-like phenotype, found reduced ENCC migration speed was observed when a subset of ENCCs were imaged during a window of time during colonization; however, proliferation was not noted to play a role in ENS phenotypes (Heanue et al., 2016). It should be noted that Heanue et al. utilized BrdU staining to assay proliferation, which labels S-phase of the cell cycle rather than M-phase labeling with pHH3. Additionally, Heanue et al. assayed the single 48 hpf timepoint, potentially missing changes in ENCC proliferation present within their *ret*^*+/-*^ zebrafish model that we observed at 60 hpf. As such, incongruencies between our similar HSCR-like *ret*^*+/-*^ zebrafish models are likely attributed to differences in assays performed, as well as timepoints assayed.

Imaging *in toto* gut colonization enabled us to spatiotemporally map the location of each ENCC generation throughout the full migratory phase of ENS formation *in vivo*, an undertaking not achieved in previous studies. From our study, we observed that vast generations of ENCCs colonize the gut. Specifically, we noticed that cell divisions, whose axis was perpendicular to the A-P axis of the gut tube, produced a growing number of ENCC generations over time, which would colonize increasingly distant domains within the gut with each subsequent generation. Our findings on global ENCC tracing adds to previous focal lineage tracing experiments performed in zebrafish, between 3.5-4 dpf (Kuwata et al., 2019). At those specific times, Kuwata and researchers found that ENCC proliferation occurred along all regions of the gut and corresponded with cell separation and displacement. Additionally, Harrison et al. found that between 50-57 hpf ENCCs in zebrafish *med24* morphants migrated along the gut tube much slower than controls, a finding that was associated with decreased mitotic rate (Harrison et al., 2014).

Collectively with prior work, our study supports the model of cell number dependent migration, whereby proliferation is required to maintain optimal inter-cell distances among the population of migrating ENCCs in order to sustain the posterior expansion, an observation congruent to the “frontal expansion” model of NCC migration that was previous postulated following mathematical modeling (Simpson et al., 2007).

Previous work done by others as well as ourselves, found the TALE homeodomain transcription factor encoding transcript, *Pbx3*, to demarcate the transition of enteric neuroblast to differentiated enteric sensory neuron subtype, IPAN (Howard, Baker et al., 2021; Ii et al., 2020; Morarach et al., 2021). Here, we demonstrate that *ret* heterozygous mutant background is capable of inducing premature expression of *pbx3b* during ENCC migration, providing evidence that Ret may serve to regulate the timing of neuronal differentiation among enteric neuroblasts. While we observed higher levels of *pbx3b* expression in *ret*^*wmr1/+*^ ENCCs compared to *ret*^*+/+*^ controls between 72-96 hpf, the levels of *pbx3b* expression were found to be the same between *ret*^*wmr1/+*^ and *ret*^*+/+*^ fish by 120 hpf, suggesting premature neuronal differentiation rather than a change in enteric subtype specification. Collectively, these results argue that the reduction in proliferative expansion of *ret*^*wmr1/+*^ ENCCs is correlated with the timing of their neuronal differentiation.

Our findings suggest that Ret signaling may function to regulate maintenance of a stem-state in enteric neural progenitors, a function that is likely conserved within the closely related vagal-NCC derived sympathoadrenal precursors that have been shown to form neuroblastoma in the presence of constitutively active Ret mutations (Li et al., 2019). Ret inhibition has been shown to significantly reduce tumor growth in a neuroblastoma mouse model (Cazes et al., 2014), further supporting evidence that Ret signaling serves to regulate NCC-derived, neural progenitor proliferation. Mutations in *ret* have also been linked to additional human cancers including multiple endocrine neoplasia 2 (MEN2), papillary thyroid carcinoma (PTC), and non-small cell lung cancer (NSCLC) (Cazes et al., 2014; Li et al., 2019). In the case of these cancers, the *ret* mutations are found to be gain-of-function in nature, while *ret* missense and nonsense mutations found to cause HSCR are distinctly loss-of-function (Takahashi, 2001). In cancer, we see *ret* associated with tumorigenesis, while in HSCR we see a reduction in overall neuroblast number, collectively suggesting *ret* functions to regulate cell number.

Recently, a functional link between Pbx3 and Meis1 has been discovered during leukemia progression where Pbx3 and Meis1 directly interact to promote the stability of Meis1, together co-driving tumor formation from transduced primary cells (Garcia-Cuellar et al., 2015). Furthermore, Garcia-Cueller et al. noted that overexpression of Pbx3 led to a dramatic increase in Meis1 transcription, suggesting genetic interactions. In the developing nervous system, Pbx and Meis factors are well-known regulators of neurogenesis, and thus, their prolonged/enhanced expression could tip the balance between proliferation and differentiation, which remains to be tested in the ENS. The findings of this study implicate Pbx3 signaling downstream of Ret and highlight an area for continued investigation.

## Methods and Materials

### Zebrafish Husbandry

All work was performed under protocols approved by, and in accordance with, the Rice University Institutional Animal Care and Use Committee (IACUC). (*Danio rerio)* Tg(*-8*.*3phox2bb*:Kaede)^em2Tg^ (Harrison et al., 2014a) adults were maintained as outcrosses with AB wildtype in our in-house zebrafish facility at 28.5 °C on a 13-hour light/11 hour dark cycle. Tg(*-8*.*3phox2bb*:Kaede);*ret*^*wmr1/+*^ fishline was maintained as AB outcrossed F1 adults, F2 embryos were collected following in-cross and processed identically to WT embryos. F2 Adults were in-crossed, in the resulting embryos the hours post fertilization (hpf) was carefully recorded, and the embryos were sorted for Kaede^+^ individuals. All embryos were cultured in standard E3 media until 24 hpf, then transferred to a 1X 1-phenyl 2-thiourea (PTU)/E3 solution (Sigma-Aldrich, #P7629) (Karlsson et al., 2001), to prevent the formation of melanin pigment.

### CRISPR-Cas9 guide design

A sgRNA targeting the 8^th^ exon of *ret* (GAGACAGCGAGGAGACTGTG) was designed by manually searching the *ret* CDS (ENSDARG00000055305.7) for protospacer adjacent motifs (PAM) within coding sequence (CDS) domains. The generation of sgRNA and generation of *cas9* mRNA were based on previously described work (Clements et al., 2017; Gagnon et al., 2014). Briefly, these methods included the use of MEGAscript *in vitro* transcription kit (Invitrogen, AM1330) to generate *cas9* mRNA from a pCS2-nls-cas9 vector (Jao et al., 2013) and the sgRNA from custom DNA oligo containing a SP6 promoter, the details previously described (Gagnon et al., 2014).

### CRISPR-Cas9 microinjection, *ret*^*wmr1/+*^ fishline establishment and genotyping, *ret*^*wmr1*^ mRNA functional analysis

Zebrafish embryos obtained from in-crossing Tg(−8.3*phox2bb*:Kaede)^+/-^ adults were injected through the chorion at the 1-cell stage with a cocktail containing phenol red dye (1:8 *µ*l), 150 picograms (pg) *cas9* mRNA and 40 pg sgRNA targeting the 8^th^ exon of *ret* (GAGACAGCGAGGAGACTGTG) (as described above). Injected F0 embryos were raised to 5 dpf, and screened for ENS defects in which regions of the intestine lacked Kaede^+^ cells in order to validate successful CRISPR-Cas9 activity (**Fig. S4**). Injected embryos exhibiting WT phenotype were collected and raised in our in-house zebrafish facility, and when reaching sexual maturity were outcrossed to WT AB. Resulting F1 embryos were screened for WT phenotype at 5 dpf and raised to sexual maturity. Resulting F1 adults were fin clipped and genotyped (Meeker et al., 2007) using forward (GCTATGCGGAATGCAATAGC) and reverse (AATCCTGAGGACAGATGGAG) primers to produce 561 bp amplicon from the *ret* locus. F1 heterozygous adults harboring identical 11 bp deletions (as described in results) in *ret* were designated *ret*^*wmr1/+*^ fishline. Functional loss of *ret*^*wmr1*^ was validated by injecting 30 pg of *ret9*^*wmr1*^ and *ret9*^*WT*^ mRNA into F2 embryos obtained from an incross of the *ret*^*wmr1/+*^ fishline. The ret9 isoform has previously been shown to be sufficient to drive ENS formation in zebrafish (Heanue and Pachnis, 2008). Injected F2 embryos were screened at 96 hpf and scored for total number of embryos exhibiting either: WT (*ret*^*+/+*^), hypoganglionosis (HSCR-like) (*ret*^*wmr1/+*^), or total aganglionosis (*ret*^*wmr1/wmr1*^) phenotypes (**Fig. S2**). In order to generate *ret9*^*wmr1*^ and *ret9*^*WT*^ mRNA, *ret*^*+/+*^ and *ret*^*wmr1/wmr1*^ embryos were separately processed for total RNA extraction using standard Trizol/Chloroform RNA precipitation. Total RNA extractions were used to generate *ret*^*+/+*^ and *ret*^*wmr1/wmr1*^ cDNA libraries using SuperScript IV kit (Invitrogen: REF 18091050). Both *ret9* (ENSDART00000139237.3) and *ret51* (ENSDART00000077627.7) splice variants were amplified from cDNA libraries using forward (GGCTCCTTTCGCTCGAATCA) and reverse (ACACTCAGCTTAATGTAGTTATTGTTGCAC) primers and Phusion PCR (Invitrogen). *ret9* and *ret51* splice variant amplicons were cloned into pCR-Blunt II-TOPO vector using manufactures instructions (Invitrogen: REF 45-0245). Zero Blunt TOPO reactions were transformed into Max Efficiency DH5α bacteria (Invitrogen: REF 18258-012). Bacteria were mini-prepped and screened using restriction digest cloning. Presumptive plasmids containing *ret9*^*wmr1*^ and *ret9*^*WT*^ cDNA were confirmed following whole-plasmid sequencing by Plasmidsaurus (https://www.plasmidsaurus.com). PolyA sequences were amplified from pCS2+ vector and fastcloned (Li et al., 2011) into TOPO-*ret9*^*wmr1*^ and TOPO-*ret9*^*WT*^ plasmids using forward (PolyA: CCTTCCATAGGAAGAGCTGTGATCCAGACATGATAAGATAC) (TOPO: CAAGCTTGATGCATAGCTTGAG) and reverse (PolyA: CAAGCTATGCATCAAGCTTGGTTAACTTGTTTATTGCAGC) (TOPO: ACAGCTCTTCCTATGGAAGG) primers. Fastcloning reactions were transformed, prepped, and validated using previously mentioned methods. Sequenced validated TOPO-*ret9*^*wmr1*^-PolyA and TOPO-*ret9*^*WT*^-PolyA plasmids were linearized with HpaI restriction enzyme and used to generate *ret9*^*wmr1*^ and *ret9*^*WT*^ mRNA using mMESSAGE mMACHINE T7 *in vitro* mRNA synthesis (Invitrogen: REF AM1344). mRNA synthesis reactions were cleaned using Monarch RNA Cleanup Kit (NEB: #T2040L) and visualized using agarose gel-electrophoresis. Pure *ret9*^*wmr1*^ and *ret9*^*WT*^ mRNA was titrated and quantified using nanodrop to inject 30 pg using same methods described for CRISPR-Cas9 injections.

### Confocal Time-Lapse microscopy of *In Toto* ENCC migration

F2 embryos from an in-cross of F1 Tg(*-8*.*3phox2bb*:Kaede);*ret*^*wmr1/+*^ fish were manually dechorionated using fine forceps, then mounted in 15 *µ*-Slide 4 Well imaging chambers (Ibidi, 80427) using 1.0% low melt temperature agarose dissolved in 1X PTU/E3 media. Embedded embryos were then covered in E3 media supplemented with 0.4% Tricane (Sigma, A5040) and 1X PTU/E3 media. Confocal time-lapse microscopy was performed with an Olympus FV3000 and FluoView software (2.4.1.198), using a long working distance 20.0X objective (UCPLFLN20X) at a constant temperature of 28°C using OKOLAB Uno-controller imaging incubator. 4D time-lapse was performed to capture full gut (*in toto*) volumes along the X-Y-Z planes by using a tiling method to capture the full-length of the gut in 2 ROIs, at ∼524-1314 second intervals for 48 consecutive hours, covering 48-96 hpf of development. *ret*^*+/+*^ (N=3), *ret*^*wmr1/+*^ (N=3). Time lapse datasets were stitched in Cellsens software and were exported and saved as .oir files, then processed further using IMARIS, as described below. Fidelity of cell tracking to discriminate individual ENCCs was ensured using a H2A-mCherry marker in alternative fishline, Tg(*-8*.*3phox2bb*:Kaede;H2A-mCherry; *unpublished*) (**Fig. S5**).

### Enteric cell tracking and generation analysis

Cell tracking was performed using spots lineage-tracking function in IMARIS (version 9.7.2). The position of every individual ENCC body (spot) was tracked across each time point along the X-Y-Z planes using manual spot detection in the Kaede channel to determine the migratory track and generation of the entire ENCC population per fish (**Fig. S1**). Every ENCC was labeled manually within the spot lineage tracks by applying a unique ID to denote cell generation as a decimal. Generation ID’s were empirically assigned based on assumed asymmetrical division along the anterior-posterior axis. That is, as a tracked cell gave rise to two cells by dividing along an axis perpendicular to the A-P axis of the gut tube, the anterior daughter cell retained the ID of the parent cell, while the posterior daughter obtained a new generation ID whose decimal denoted +1 the parent cell. Generations were viewed in the Vantage viewer of IMARIS for global visualizations. Vanguard cells were defined as the leading-edge migratory cells along the gut. Therefore, the leader identity (Vanguard) may change when the current leader is overtaken by a new cell born by cell division during migration. Total raw data of spots tracking was exported from IMARIS and further analyzed for graphical depictions using R studio (Version 1.3.959) [Package *ggplot2* version 3.3.2]. For data depicting cell “Distance from Foregut,” distance was defined based on origin reference frames placed in the anterior foregut, at the level of 13-14^th^ somite anterior to the cloaca.

### Hybridization chain reaction and phospho-histone H3 whole-mount immunohistochemistry

Hybridization chain reaction experiments were performed in accordance with previously described methods (Howard, Baker et al., 2021; Ibarra-García-Padilla et al., 2021), Ref Seq IDs *vipb* (NM_001114555.1) *pbx3b* (BC131865.1). Whole-mount immunofluorescence experiments were conducted according to methods previously described (Baker et al., 2019). All F2 *ret*^*wmr1/+*^ embryos used for fixed tissue experiments were screened for the HSCR-like phenotype in which Kaede^+^ ENCCs had failed to reach the hindgut at 72 hpf, 80 hpf, 96 hpf, and 120 hpf. Experiments using 60 hpf fixed tissue were not able to be sorted based on phenotype. For these experiments, individual F2 embryos obtained from F1 *ret*^*wmr1/+*^ in-cross were fixed in individual wells within a 96 well plate following careful dissection of the head. Head tissue was processed for genomic DNA (gDNA) in separate 96 well plates with 30 *µ*l NaOH. Heads were boiled at 95°C for 10 min, briefly vortexed, then pH balanced with 3 *µ*l 1M Tris HCL pH 8.0 and stored at -20°C. gDNA was used as template for PCR using forward (GCCACGATTTCCAGCTTGTGC) and reverse (ACTGGCATCTCCCTGTTGG) primers that amplify a small, 89 bp segment surrounding the lesion site. PCR product was processed using T7 endonuclease assay **(Fig. S6)** per manufacturers instruction (NEB #E3321) to identify *ret*^*wmr1/+*^ embryos which were then pooled and processed via previously described HCR and whole-mount immunohistochemical preparations. The primary antibodies were used at the following dilutions: Mouse monoclonal anti-phospho-Histone H3 IgG1 (Abcam ab14955) 1:1500, rabbit polyclonal anti-Kaede IgG (MBL international PM102M) 1:250, and mouse monoclonal anti-HuC/HuD (Elavl3/4) IgG2b (Invitrogen Molecular Probes A21271) 1:200. The secondary antibodies were ordered from Invitrogen and used at the following dilutions: Alexa Fluor 488 Goat anti-Rabbit IgG (A11008) 1:500, Alexa Fluor 647 Goat anti-Mouse IgG2b (A21242) 1:500, and Alexa Fluor 568 Goat anti-Mouse IgG1b (A21124) 1:500. Confocal microscopy was performed with an Olympus FV3000 and FluoView software (2.4.1.198), using a long working distance 20.0X objective (UCPLFLN20X).

### Statistics

Statistical analysis was performed in R studio software (Version 1.3.959) using [Package *ggsignif* version 0.6.0] and [Package *stats* version 4.0.2]. Shapiro-Wilk was used to test normality. For comparisons, normally distributed data was tested using unpaired *t* Test and non-normally distributed using Wilcoxon signed-rank Test. *P* < 0.05 (NS = non-significant) *P* < 0.05 (*), *P* < 0.01 (**), and *P* < 0.001 (***). Post *hoc* power analysis was performed using G*Power 3 and found sufficient statistical power (>0.8) for all tests in which we report statistical significance (Faul et al., 2007).

## Acknowledgements

We acknowledge funding of this project from Rice University, Cancer Prevention and Research Institute of Texas (CPRIT) Recruitment of First-Time Tenure Track Faculty Members (CPRIT-RR170062) to R.A.U., from NSF CAREER Award (1942019) awarded to R.A.U., and National Institutes of Health DK124804 awarded to R.A.U. IMARIS image analysis was performed using Rice University’s Shared Equipment Authority (SEA) IMARIS workstation and we thank Alloysius Budi Utama for his insights and training with this software. We thank Dan Wagner (Rice University) for helpful advice throughout this project. We also thank Thom Yorke and Simon Green for technical assistance.

## Figures

**Figure S1.**
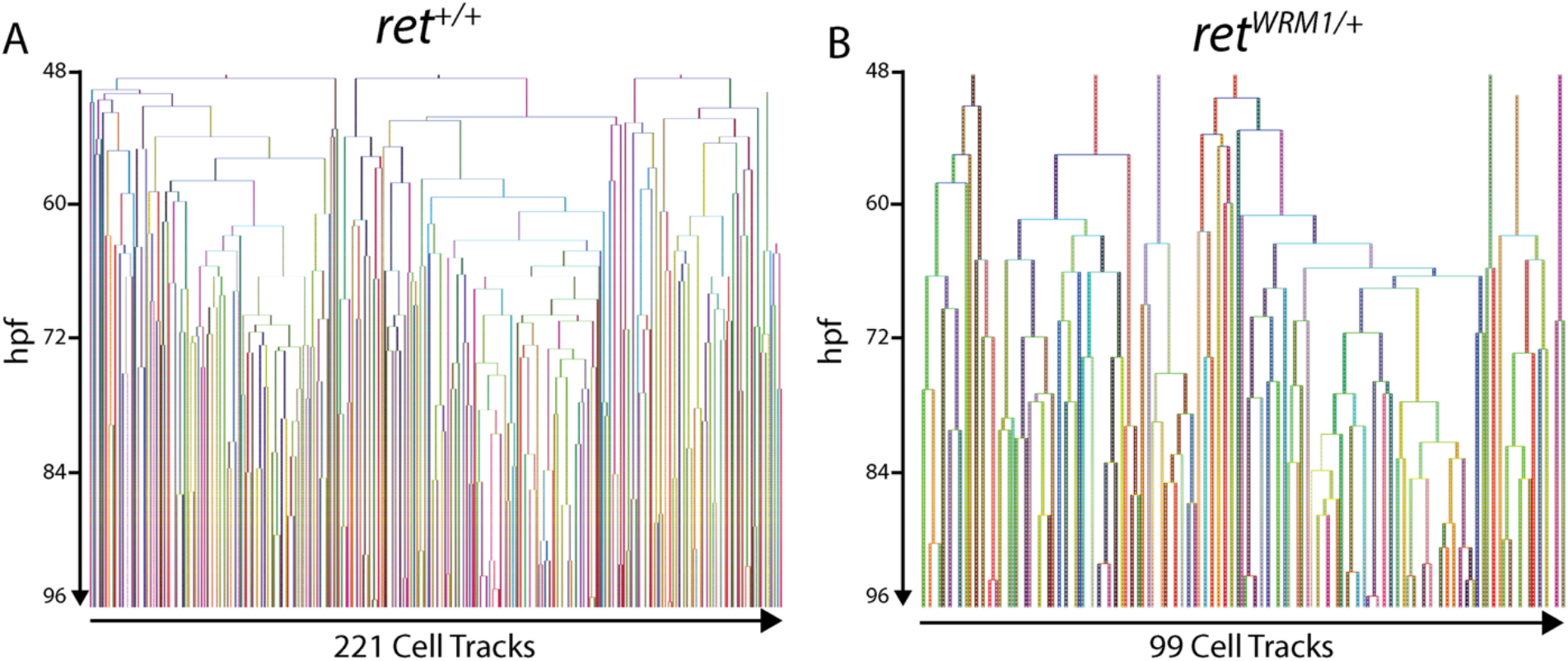
IMARIS snapshots of ENCC generation trees from representative 48-96 hpf time-lapses. (A) *ret*^*+/+*^ ENCC generation tree reveals emergence of 221 ENCCs between 48-96 hpf. (B) *ret*^*wmr1/+*^ ENCC generation tree reveals emergence of 99 ENCCs between 48-96 hpf.

**Figure S2.**
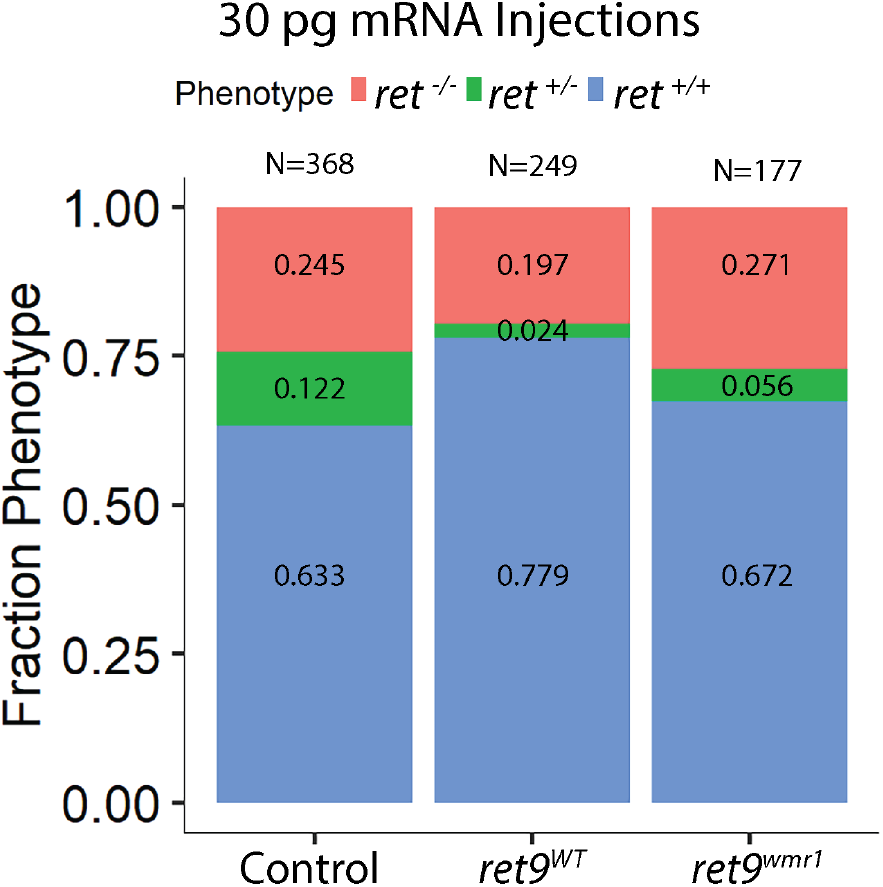
*ret*^*WT*^ mRNA partially rescues the percentage of HSCR phenotypes seen in larvae from a *ret*^*wmr1/+*^ incross. Stacked bar graphs depict fraction of *ret*^*+/+*^ (WT; Blue), *ret*^*+/-*^ (HSCR-like; Green) and *ret*^*-/-*^ (Total Agangionosis; Magenta) phenotypes scored in *ret*^*wmr1/+*^ incross F2’s at 96 hpf following uninjected control, 30 pg *ret9*^*WT*^ mRNA and 30 pg *ret9*^*wmr1*^ mRNA. *ret9*^*WT*^ mRNA shows partial rescue of disease phenotypes while *ret9*^*wmr1*^ mRNA fails to rescue, demonstrating loss-of-function.

**Figure S3.**
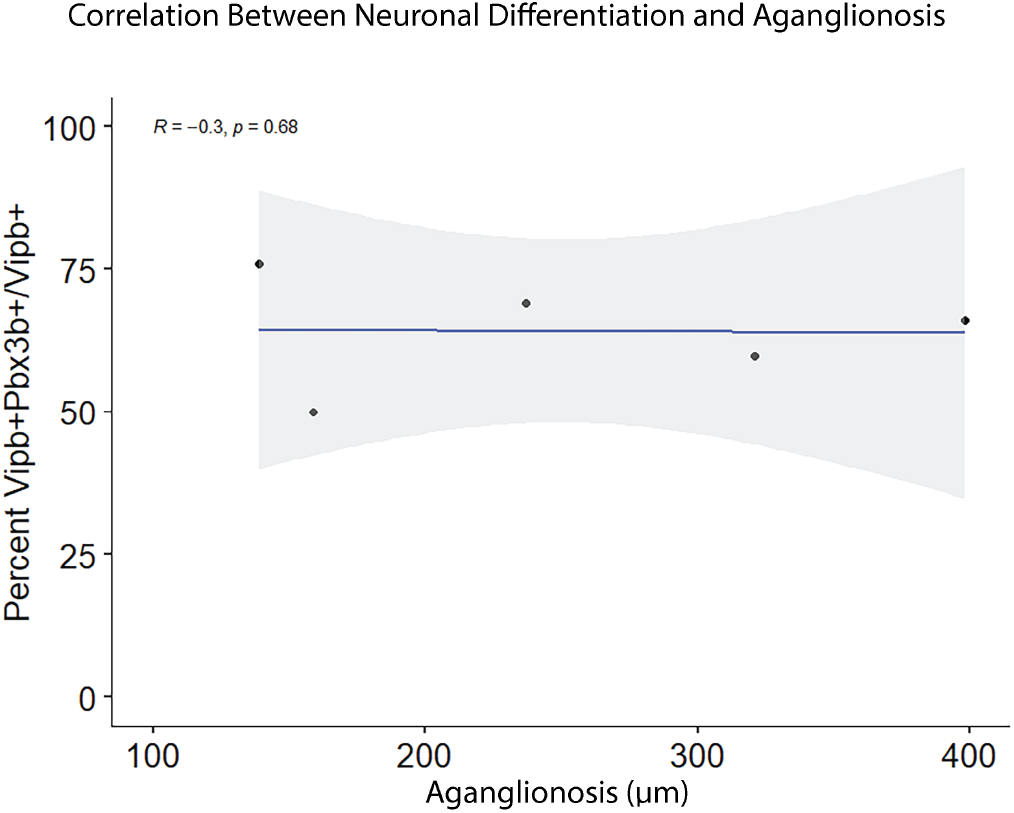
No correlation between *pbx3b*^*+*^ enteric neuronal differentiation and extent of aganglionosis in *ret*^*wmr1/+*^ larvae at 96 hpf. *ret*^*wmr1/+*^ larvae used in figure 6G were analyzed to measure the distance between distal most *vipb*^*+*^ ENCCs and cloaca (Aganglionosis (*µ*m)). Scatter plot depicting percent *pbx3b*^*+*^ (*vipb*^*+*^*pbx3b*^*+*^*/vipb*^*+*^) vs extent of aganglionosis (*µ*m) tested using Spearman correlation (*R=*-0.3) finds no correlation between the two variables.

**Figure S4.**
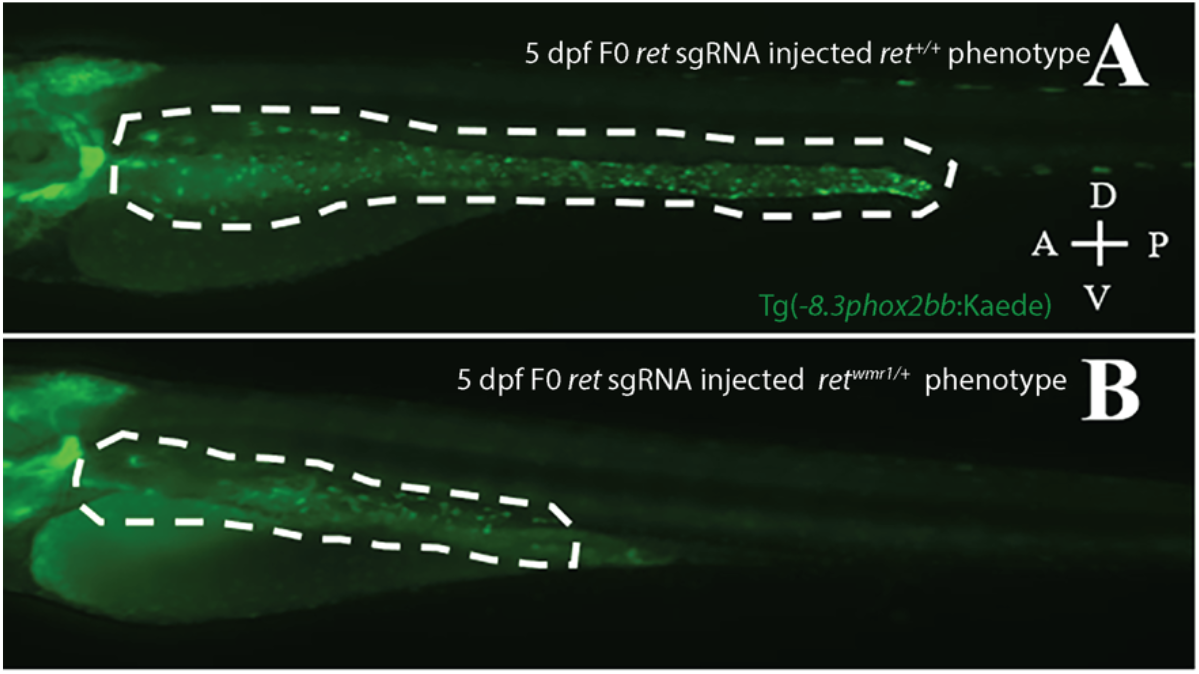
Phenotypic crispant screen of F0 *ret* sgRNA injected Tg(*-8*.*3phox2bb:*Kaede) larvae. (A) 5 dpf *ret* sgRNA injected Tg(*-8*.*3phox2bb:*Kaede) larvae exhibiting *ret*^*+/+*^ phenotype with Kaede^+^ ENCCs throughout the gut. (B) 5 dpf *ret* sgRNA injected Tg(*-8*.*3phox2bb:*Kaede) larvae exhibiting *ret*^*wmr1/+*^ phenotype lacking Kaede^+^ ENCCs throughout the hindgut. Gut tube colonized by ENCCs outlined by white dashed lines. A: anterior; P: posterior; D: dorsal; V: ventral, shown in A.

**Figure S5.**
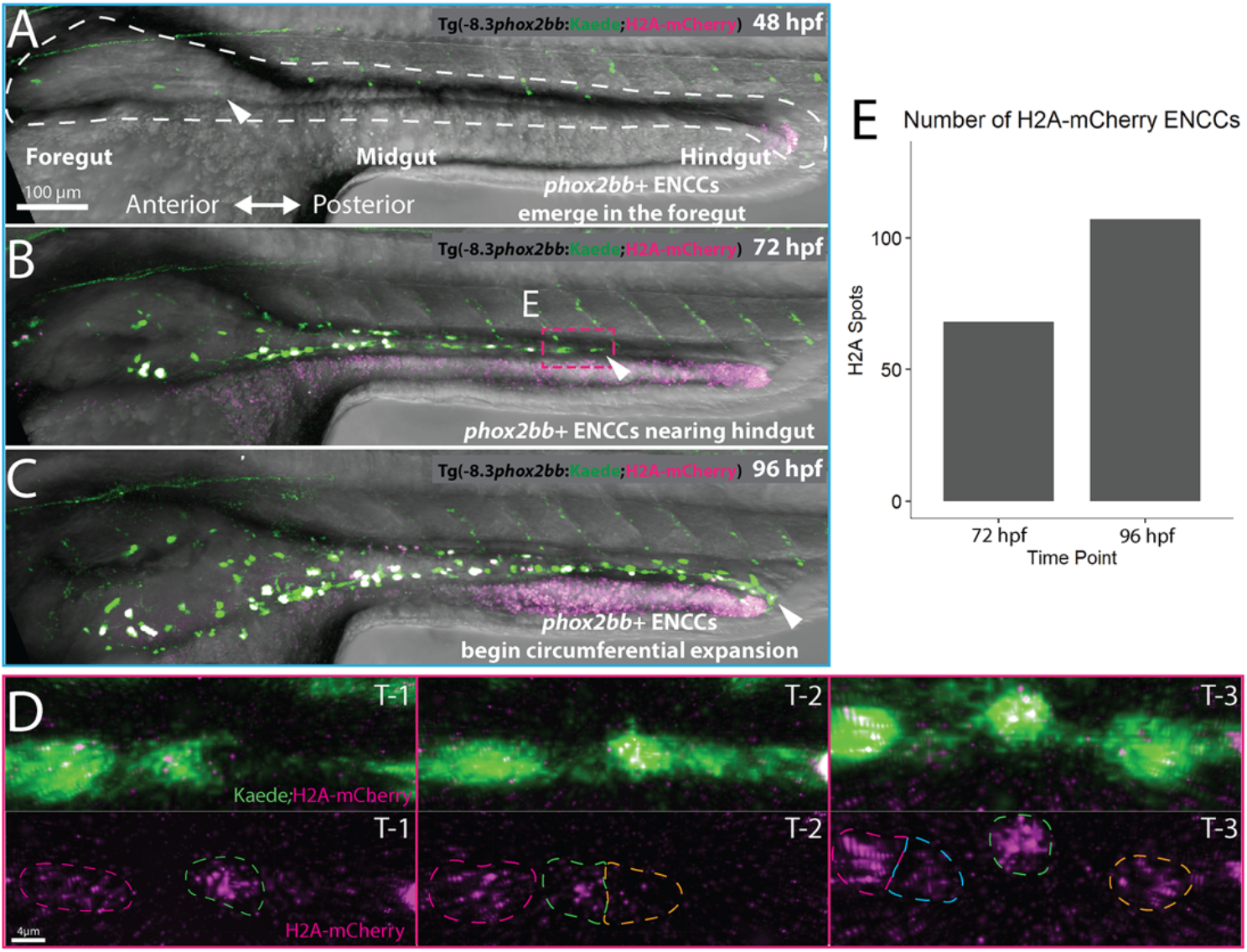
4D time-lapse microscopy between 48-96 hpf in transgenic line Tg(−8.3*phox2bb*:Kaede;H2A-mCherry). (A-C) Still images from time-lapse reveal (A, white arrow) *phox2bb*^*+*^ ENCCs emerging in foregut intestine at 48hpf (B, white arrow), leading ENCCs reaching the hindgut at 72 hpf (C), and the circumferential expansion of ENCCs at 96 hpf. (D) Expanded images of cell migration and division at the ENCC wavefront seen in panel B, magenta box. Time-points 1-3 (T1-3) depict representative time-points where cell divisions occur. (E) Quantifications of H2A-mCherry+ cells at 72 and 96 hpf reveal comparable ENCCs number to *ret*^*wmr1/+*^ *phox2bb:*Kaede cell counts (Fig. 3A).

**Figure S6.**
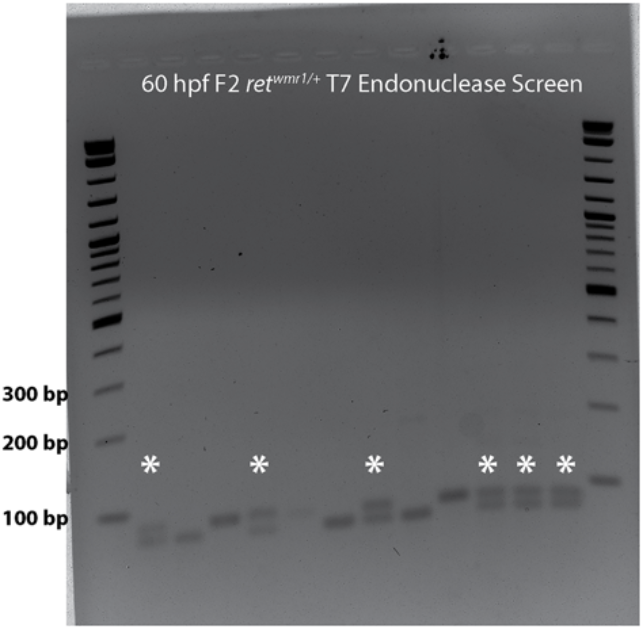
T7 endonuclease assay identifies *ret*^*wmr1/+*^ genotype in 60 hpf F2 larvae. 3% agarose gel used to separate T7 endonuclease cleaved PCR product of *ret* loci identifies heterozygotic *ret*^*wmr1/+*^ larvae (starred lanes) from genomic DNA isolated from dissected heads. Heterozygotic larvae collected and pooled for fixed tissue hybridization chain reaction and whole-mount immunofluorescence assays.

**Table S1.** Phenotypic scores of 5 dpf F2 embryos obtained from *ret*^*wmr1/+*^ in-cross. Table shows total counts and percent prevalence of *ret*^*+/+*^ (WT), *ret*^*wmr1/+*^ (HSCR-like), and *ret*^*wm1/wmr1*^ (Total aganglionosis) embryos.

| Plate |  | F2 Ret<sup>WMR1/+</sup> |
| --- | --- | --- |
| "WT Phenotype" (+/+) | Counts | 32 |
|  | Percent | 16.93 |
| HSCR Phenotype (-/+) | Counts | 53 |
|  | Percent | 28.04 |
| No Cells in ENS (-/-) | Counts | 104 |
|  | Percent | 55.03 |
| Total |  | 189 |

## Movie Legends

**Movie 1**. Enteric neural crest cells migrate posteriorly down a control zebrafish larval gut between 48-96 hpf. Lateral view of *phox2bb:*Kaede^+^ enteric neural crest cells appear in the foregut posterior to origin reference frame and continue to propagate in number as they migrate posteriorly and colonize the entire length of the gut tube.

**Movie 2**. Enteric neural crest cells migrate posteriorly down a control zebrafish larval gut between 48-96 hpf, with IMARIS spots shown. Same movie as Movie 1 with cell spots overlaid with each individual *phox2bb:*Kaede^+^ enteric neural crest cell (color coded based on unique ID). Large turquoise and pink spots correspond to leading most (vanguard) cell in the right and left migratory cell chains, respectively. Tracks correspond to vanguard migratory track and are color coded based on time during movie.

**Movie 3**. Animated 360° horizontal rotation of cell tracks from Figure 1D and Movie 2. Tracks color coded cold-hot for timepoints throughout time-lapse.

**Movie 4**. Animated 360° vertical rotation of cell tracks from Figure 1D and Movie 2. Tracks color coded cold-hot for timepoints throughout time-lapse.

**Movie 5**. Enteric neural crest cells migrate posteriorly down *ret*^*wmr1/+*^ zebrafish larval gut between 48-96 hpf. Lateral view of *phox2bb:*Kaede^+^ enteric neural crest cells appear in the foregut posterior to origin reference frame and migrate posteriorly yet fail to colonize the entire length of the gut tube.

**Movie 6**. Enteric neural crest cells migrate posteriorly down *ret*^*wmr1/+*^ zebrafish larval gut between 48-96 hpf, with IMARIS spots shown. Same movie as Movie 3 with cell spots overlaid with each individual *phox2bb:*Kaede^+^ enteric neural crest cell (color coded based on unique ID). Large turquoise and pink spots correspond to leading most (vanguard) cell in the right and left migratory cell chains, respectively. Tracks correspond to vanguard migratory track and are color coded based on time during movie.

## References

Amiel, J., Sproat-Emison, E., Garcia-Barcelo, M., Lantieri, F., Burzynski, G., Borrego, S., Pelet, A., Arnold, S., Miao, X., Griseri, P., et al. (2008). Hirschsprung disease, associated syndromes and genetics: A review. J. Med. Genet. 45, 1–14.

Baker, P.A., Meyer, M.D., Tsang, A., and Uribe, R.A. (2019). Immunohistochemical and ultrastructural analysis of the maturing larval zebrafish enteric nervous system reveals the formation of a neuropil pattern. Sci. Rep. 9, 6941.

Barlow, A.J., Wallace, A.S., Thapar, N., and Burns, A.J. (2008). Critical numbers of neural crest cells are required in the pathways from the neural tube to the foregut to ensure complete enteric nervous system formation. Development 135, 1681–1691.

Cazes, A., Lopez-Delisle, L., Tsarovina, K., Pierre-Eugène, C., De Preter, K., Peuchmaur, M., Nicolas, A., Provost, C., Louis-Brennetot, C., Daveau, R., et al. (2014). Activated Alk triggers prolonged neurogenesis and Ret upregulation providing a therapeutic target in ALK-mutated neuroblastoma.

Clements, T.P., Tandon, B., Lintel, H.A., McCarty, J.H., and Wagner, D.S. (2017). RICE CRISPR: Rapidly increased cut ends by an exonuclease Cas9 fusion in zebrafish. Genesis 55, 1–6.

Faul, F., Erdfelder, E., Lang, A.G., and Buchner, A. (2007). G*Power 3: A flexible statistical power analysis program for the social, behavioral, and biomedical sciences. Behav. Res. Methods 2007 392 39, 175–191.

Furness, J.B., and Wiley InterScience (Online service) (2006). The enteric nervous system (Blackwell Pub).

Furness, J.B., Jones, C., Nurgali, K., and Clerc, N. (2004). Intrinsic primary afferent neurons and nerve circuits within the intestine. Prog. Neurobiol. 72, 143–164.

Gagnon, J.A., Valen, E., Thyme, S.B., Huang, P., Ahkmetova, L., Pauli, A., Montague, T.G., Zimmerman, S., Richter, C., and Schier, A.F. (2014). Efficient Mutagenesis by Cas9 Protein-Mediated Oligonucleotide Insertion and Large-Scale Assessment of Single-Guide RNAs. PLoS One 9, e98186.

Ganz, J. (2018). Gut feelings: Studying enteric nervous system development, function, and disease in the zebrafish model system. Dev. Dyn. 247, 268–278.

Garcia-Cuellar, M.P., Steger, J., Füller, E., Hetzner, K., and Slany, R.K. (2015). Pbx3 and Meis1 cooperate through multiple mechanisms to support Hox-induced murine leukemia. Haematologica 100, 905.

Gurley, L.R., D’Anna, J.A., Barham, S.S., Deaven, L.L., and Tobey, R.A. (1978). Histone Phosphorylation and Chromatin Structure during Mitosis in Chinese Hamster Cells. Eur. J. Biochem. 84, 1–15.

Harris, M.L., and Erickson, C.A. (2007). Lineage specification in neural crest cell pathfinding. Dev. Dyn. 236, 1–19.

Harrison, C., Wabbersen, T., and Shepherd, I.T. (2014). In vivo visualization of the development of the enteric nervous system using a Tg(−8.3bphox2b:Kaede) transgenic zebrafish. Genesis 52, 985–990.

Heanue, T.A., and Pachnis, V. (2007). Enteric nervous system development and Hirschsprung’s disease: Advances in genetic and stem cell studies. Nat. Rev. Neurosci. 8, 466–479.

Heanue T.A., Pachnis V. (2008). Ret isoform function and marker gene expression in the enteric nervous system is conserved across diverse vertebrate species. Mech Dev. Aug;125(8):687–99.

Heanue, T.A., Boesmans, W., Bell, D.M., Kawakami, K., Vanden Berghe, P., and Pachnis, V. (2016). A Novel Zebrafish ret Heterozygous Model of Hirschsprung Disease Identifies a Functional Role for mapk10 as a Modifier of Enteric Nervous System Phenotype Severity. PLOS Genet. 12, e1006439.

Hendzel, M.J., Wei, Y., Mancini, M.A., Van Hooser, A., Ranalli, T., Brinkley, B.R., Bazett-Jones, D.P., and Allis, C.D. (1997). Mitosis-specific phosphorylation of histone H3 initiates primarily within pericentromeric heterochromatin during G2 and spreads in an ordered fashion coincident with mitotic chromosome condensation. Chromosom. 1997 1066 106, 348–360.

Hockley, J.R.F., Taylor, T.S., Callejo, G., Wilbrey, A.L., Gutteridge, A., Bach, K., Winchester, W.J., Bulmer, D.C., McMurray, G., and Smith, E.S.J. (2019). Single-cell RNAseq reveals seven classes of colonic sensory neuron. Gut 68, 633–644.

Howard, A.G., Baker, P.A., Ibarra-García-Padilla, R., Moore, J.A., Rivas, L.J., Tallman, J.J., Singleton, E.W., Westheimer, J.L., Corteguera, J.A., and Uribe, R.A. (2021). An atlas of neural crest lineages along the posterior developing zebrafish at single-cell resolution. Elife 10.

Howard, A.G.A., Nguyen, A.C., Tworig, J., Ravisankar, P., Singleton, E.W., Li, C., Kotzur, G., Waxman, J.S., and Uribe, R.A. (2022). Elevated Hoxb5b Expands Vagal Neural Crest Pool and Blocks Enteric Neuronal Development in Zebrafish. Front. Cell Dev. Biol. 9, 1–18.

Hutchins, E.J., Kunttas, E., Piacentino, M.L., Howard, A.G.A., Bronner, M.E., and Uribe, R.A. (2018). Migration and diversification of the vagal neural crest. Dev. Biol.

Ibarra-García-Padilla, R., Howard, A.G.A., Singleton, E.W., and Uribe, R.A. (2021). A protocol for whole-mount immuno-coupled hybridization chain reaction (WICHCR) in zebrafish embryos and larvae. STAR Protoc. 2, 100709.

Ii, M.A.F., Ehsan, L., Moore, S.R., and Levin, D.E. (2020). The Enteric Nervous System and Its Emerging Role as a Therapeutic Target.

Jao, L.E., Wente, S.R., and Chen, W. (2013). Efficient multiplex biallelic zebrafish genome editing using a CRISPR nuclease system. Proc. Natl. Acad. Sci. U. S. A. 110, 13904–13909.

Karlsson, J., Von Hofsten, J., and Olsson, E. (2001). Generating Transparent Zebrafish: A Refined Method to Improve Detection of Gene Expression During Embryonic Development.

Khomgrit, M., Anastassia, M., Viktoria, K., Fatima, M., Rakesh, K., Wei, L., Patrik, E., Ulrika, M., Marklund, U., and Institute, K. Diversification of molecularly defined myenteric neuron classes revealed by single cell RNA-sequencing.

Kucenas, S., Takada, N., Park, H.C., Woodruff, E., Broadie, K., and Appel, B. (2008). CNS-derived glia ensheath peripheral nerves and mediate motor root development. Nat. Neurosci. 11, 143–151.

Kuo, B.R., and Erickson, C.A. (2011). Vagal neural crest cell migratory behavior: A transition between the cranial and trunk crest. Dev. Dyn. 240, 2084–2100.

Kuwata, M., Nikaido, M., and Hatta, K. (2019). Local heat-shock mediated multi-color labeling visualizing behaviors of enteric neural crest cells associated with division and neurogenesis in zebrafish gut. Dev. Dyn. 248, 437–448.

Lake, J.I., and Heuckeroth, R.O. (2013). Enteric nervous system development: Migration, differentiation, and disease. Am. J. Physiol. - Gastrointest. Liver Physiol. 305, G1.

Landman, K.A., Simpson, M.J., and Newgreen, D.F. (2007). Mathematical and experimental insights into the development of the enteric nervous system and Hirschsprung’s Disease. Dev. Growth Differ 49, 277–286.

Li, A.Y., McCusker, M.G., Russo, A., Scilla, K.A., Gittens, A., Arensmeyer, K., Mehra, R., Adamo, V., and Rolfo, C. (2019). RET fusions in solid tumors. Cancer Treat. Rev. 81.

Li, C., Wen, A., Shen, B., Lu, J., Huang, Y., and Chang, Y. (2011). FastCloning: A highly simplified, purification-free, sequence- and ligation-independent PCR cloning method. BMC Biotechnol. 11, 92.

Meeker, N.D., Hutchinson, S.A., Ho, L., and Trede, N.S. (2007). Method for isolation of PCR-ready genomic DNA from zebrafish tissues. Biotechniques 43, 610–614.

Morarach, K., Mikhailova, A., Knoflach, V., Memic, F., Kumar, R., Li, W., Ernfors, P., and Marklund, U. (2021). Diversification of molecularly defined myenteric neuron classes revealed by single-cell RNA sequencing. Nat. Neurosci. 24, 34–46.

Nagy, N., and Goldstein, A.M. (2017). Enteric nervous system development: A crest cell’s journey from neural tube to colon. Semin. Cell Dev. Biol. 66, 94–106.

Natarajan, D., Marcos-Gutierrez, C., Pachnis, V., and de Graaff, E. (2002). Requirement of signalling by receptor tyrosine kinase RET for the directed migration of enteric nervous system progenitor cells during mammalian embryogenesis. Development 129, 5151–5160.

Natarajan, D., McCann, C., Dattani, J., Pachnis, V., Thapar, N. (2022). Multiple Roles of Ret Signalling During Enteric Neurogenesis. Front Mol Neurosci. May 27;15:832317. doi: 10.3389/fnmol.2022.832317.

Nikaido, M., Izumi, S., Ohnuki, H., Takigawa, Y., Yamasu, K., and Hatta, K. (2018). Early development of the enteric nervous system visualized by using a new transgenic zebrafish line harboring a regulatory region for choline acetyltransferase a (chata) gene. Gene Expr. Patterns 28, 12–21.

Olden, T., Akhtar, T., Beckman, S.A., and Wallace, K.N. (2008). Differentiation of the zebrafish enteric nervous system and intestinal smooth muscle. Genesis 46, 484–498.

Olsson, C., Holmberg, A., and Holmgren, S. (2008). Development of enteric and vagal innervation of the zebrafish (Danio rerio) gut. J. Comp. Neurol. 508, 756–770.

Pérez-Cadahía, B.P.-C., Drobic, B.D., and Davie, J.R.D.R. (2009). H3 phosphorylation: dual role in mitosis and interphaseThis paper is one of a selection of papers published in this Special Issue entitled 30th Annual International Asilomar Chromatin and Chromosomes Conference and has undergone the Journal’s usual peer review process. https://Doi.Org/10.1139/O09-053 87, 695–709.

Peters-Van Der Sanden, M.J.H., Kirby, M.L., Gittenberger-De Groot, A., Tibboel, D., Mulder, M.P., and Meijers, C. (1993). Ablation of Various Regions Within the Avian Vagal Neural Crest Has Differential Effects on Ganglion Formation in the Fore-, Mid-and Hindgut.

Rao, M., and Gershon, M.D. (2018). Enteric nervous system development: what could possibly go wrong? Nat. Rev. Neurosci. 19, 552–565.

Shepherd, I., and Eisen, J. (2011). Development of the zebrafish enteric nervous system. Methods Cell Biol. 101, 143–160.

Shepherd, I.T., Pietsch, J., Elworthy, S., Kelsh, R.N., and Raible, D.W. (2004). Roles for GFRα1 receptors in zebrafish enteric nervous system development. Development 131, 241–249.

Simpson, M.J., Zhang, D.C., Mariani, M., Landman, K.A., and Newgreen, D.F. (2007). Cell proliferation drives neural crest cell invasion of the intestine. Dev. Biol. 302, 553–568.

Stanchina, L., Baral, V., Robert, F., Pingault, V., Lemort, N., Pachnis, V., Goossens, M., and Bondurand, N. (2006). Interactions between Sox10, Edn3 and Ednrb during enteric nervous system and melanocyte development. Dev. Biol. 295, 232–249.

Takahashi, M. (2001). The GDNF/RET signaling pathway and human diseases. Cytokine Growth Factor Rev. 12, 361–373.

Taraviras, S. (1999). Signalling by the RET receptor tyrosine kinase and its role in the developmentof the mammalian enteric nervous system. 126, 2785–2797.

Taylor, C.R., Montagne, W.A., Eisen, J.S., and Ganz, J. (2016). Molecular Fingerprinting Delineates Progenitor Populations in the Developing Zebrafish Enteric Nervous System. Dev. Dyn. 245, 1081–1096.

Theveneau, E., and Mayor, R. (2011). Can mesenchymal cells undergo collective cell migration? The case of the neural crest. Cell Adhes. Migr. 5, 490–498.

Uesaka, T., Nagashimada, M., Yonemura, S., and Enomoto, H. (2008). Diminished Ret expression compromises neuronal survival in the colon and causes intestinal aganglionosis in mice. J. Clin. Invest. 118, 1890–1898.

Wallace, K.N., Akhter, S., Smith, E.M., Lorent, K., and Pack, M. (2005). Intestinal growth and differentiation in zebrafish. Mech. Dev. 122, 157–173.

Young, H.M., Hearn, C.J., Farlie, P.G., Canty, A.J., Thomas, P.Q., and Newgreen, D.F. (2001). GDNF is a chemoattractant for enteric neural cells. Dev. Biol. 229, 503–516.

Young, H.M., Bergner, A.J., Anderson, R.B., Enomoto, H., Milbrandt, J., Newgreen, D.F., and Whitington, P.M. (2004). Dynamics of neural crest-derived cell migration in the embryonic mouse gut. Dev. Biol. 270, 455–473.

